# Specific *SOX10* enhancer elements modulate phenotype plasticity and drug resistance in melanoma

**DOI:** 10.1101/2024.12.12.628224

**Authors:** Sophia “Noah” DeGeorgia, Charles K. Kaufman

## Abstract

Recent studies indicate that the development of drug resistance and increased invasiveness in melanoma is largely driven by transcriptional plasticity rather than canonical coding mutations. Understanding the mechanisms behind cell identity shifts in oncogenic transformation and cancer progression is crucial for advancing our understanding of melanoma and other aggressive cancers. While distinct melanoma phenotypic states have been well characterized, the processes and transcriptional controls that enable cells to shift between these states remain largely unknown. In this study, we initially leverage the well-established zebrafish melanoma model as a high-throughput system to dissect and analyze transcriptional control elements that are hijacked by melanoma. We identify key characteristics of these elements, making them translatable to human enhancer identification despite the lack of direct sequence conservation. Building on our identification of a zebrafish *sox10* enhancer necessary for melanoma initiation, we extend these findings to human melanoma, identifying two human upstream enhancer elements that are critical for full *SOX10* expression. Stable biallelic deletion of these enhancers using CRISPR-Cas9 induces a distinct phenotype shift across multiple human melanoma cell lines from a melanocytic phenotype towards an undifferentiated phenotype and is also characterized by an increase in drug resistance that mirrors clinical data including an upregulation of NTRK1, a tyrosine kinase, and potential therapeutic target. These results provide new insights into the transcriptional regulation of *SOX10* in human melanoma and underscore the role of individual enhancer elements and potentially NTRK1 in driving melanoma phenotype plasticity and drug resistance. Our work lays the groundwork for future gene-based and combination kinase-inhibitor therapies targeting *SOX10* regulation and NTRK1 as a potential avenue for enhancing the efficacy of current melanoma treatments.

## Introduction

Transcriptional regulation of cell identity is a fundamental driving force in cell biology, governing processes from embryonic development to oncogenic transformation. This regulation is particularly relevant in the context of melanoma, where oncogenic transformation involves a reversion towards developmental transcriptional programs. Melanoma, the deadliest form of skin cancer on a per case basis, arises from neural crest (NC)-derived melanocytes, which undergo oncogenic transformation via reactivation of subsets of the embryonic NC program. Notably, *SOX10*, which is critical for NC development and subsequently downregulated in mature melanocytes, is re-upregulated in melanoma cells^1–3^.

Melanoma poses a significant treatment challenge as there is frequent development of treatment resistance^4^ and a high propensity to metastasize aggressively^5^. Metastasis contributes to 90% of mortality across cancers^6^, with melanoma being no exception. Although advances in immunotherapy and targeted therapy have transformed melanoma treatment^7,8^, the inability to eradicate all residual disease allows these cancers to phenotypically adapt and metastasize while becoming treatment resistant^9^.

In various cancers, including colorectal^10^, gastric^11^, and non-small lung cancer^12^, treatment has been associated with phenotype adaptation shifts reminiscent of epithelial-mesenchymal transition (EMT)^13,14^, which has been shown to contribute to drug resistance and invasiveness. While melanoma has among the highest mutational burdens in cancer^15–17^, recent research has highlighted an additional contributor to intra-tumoral heterogeneity: transcriptional plasticity. In 2008, Hoek^18^ et al identified two primary melanoma phenotypes: proliferative and invasive. These groups were subsequently expanded to four subgroups by Tsoi et al in 2018^19^ and Rambow^9^ et al. later that year. While the nomenclature varies, most researchers agree on a continuum of phenotypic states ranging from highly differentiated, melanocytic-like states, transitioning through a neural crest-like state, and ultimately reaching a completely undifferentiated, stem cell-like state^19,20^. While the melanocyte master transcriptional regulator MITF has been a key biomarker for melanoma phenotype states, *SOX10* remains a crucial player in both melanoma cell identity both during initiation, progression, and phenotype switching^21^.

A driving force in many melanomas is the Mitogen-activated protein kinase (MAPK) pathway made up of BRAF, MEK, and ERK (leading to modulation of *MITF*, among other genes). The MAPK pathway is a signal transduction pathway that converts external stimuli to changes in gene expression^22^ and plays an important role in all eukaryotic cells, coordinating mitosis, metabolism, motility, survival, apoptosis, and differentiation^23^. Under healthy, physiological conditions, activation of the MAPK pathway leads to cell growth and proliferation. Upstream negative feedback prevents persistent MAPK pathway activation^24^. For example, MAPK-dependent p53 phosphorylation can lead to a protective halt of the cell cycle and apoptosis in some cases^25^. BRAF variants, mostly involving codon 600, occur in about 60% over melanomas, and are also in found colorectal, ovarian, and papillary thyroid carcinomas^26^. In the BRAF^V600E^ mutation (present in 70- 88% of BRAF-mutated melanomas), the valine to glutamate change in the kinase domain of the BRAF protein leads to permanent MAPK activation, regardless of negative feedback mechanisms – leading to oncogenic malignant uncontrolled growth^27^.

The first strides of progress in combatting this powerful oncogenic mechanism came with the FDA approval of BRAF and MEK inhibitor drugs between 2011 and 2013. Orally available, small-molecule drugs that selectively targeted BRAF (vemurafenib^28^ and dabrafenib^29^) or MEK (trametinib^30^ and cobimetinib^31^). Although the development of these drugs was a breakthrough in the treatment of melanoma, 15-20% of melanoma tumors harbored primary resistance to this therapy, and most responses are not durable, with most patients developing resistance to this therapy especially when presenting with a high disease burden^32^. Immunotherapies targeting PD- 1/L1, CTLA-3, and LAG-3, such as nivolumab^33^, ipilimumab^34^, and relatlimab^35^, have also considerably improved melanoma mortality, prolonging progression free survival and overall survival compared to previous treatment options, but the combinations of these therapies can be associated with significant toxicity^7,36^.

In terms of key melanoma transcriptional network regulators, targeting *SOX10* using shRNA or coding sequence deletions has shown promise in inducing cell death or phenotype shifts in melanoma^37–39^, but the transcriptional regulatory elements controlling *SOX10*’s expression in these contexts remain incompletely understood. In our recent work, we identified regulatory elements controlling *sox10* expression in zebrafish melanoma^40^ and now further use zebrafish to highlight a *sox10* enhancer relevant to melanoma onset/progression.

We then mapped these findings onto human *SOX10* regulatory regions/enhancers, focusing on conserved transcriptional motifs and enhancer features, such as the presence of a conserved *SOX10* dimer site. This cross-species approach allowed us to pinpoint two human *SOX10* enhancers, i.e. those with evolutionary conserved, paired functional *SOX10* binding sites, with likely roles in melanoma biology. We then engineered targeted deletions of these two *SOX10* enhancers and examined 13 stable enhancer deletion lines across multiple melanoma cell lines, each with varying degrees of *SOX10* dependency, to investigate how these elements impact melanoma phenotype switching. RNA-seq analysis across deletion and phenotype-switched lines revealed consistent global transcriptional shifts from melanocytic fates towards more undifferentiated fates. We also noted upregulation of specific genes like NTRK1 and other genes within the NTRK pathway that are associated with targeted therapy (i.e. BRAF/MEK inhibitor therapy) resistance in human patient samples. Knockdown of NTRK1 with siRNA and pharmacologic inhibition of NTRK1 led to increased sensitivity to BRAF and MEK inhibitors in melanoma cells. These findings highlight NTRK1 as a potential driver of drug resistance and invasiveness in melanoma in the context of loss of dependence on MITF/SOX10, suggesting a novel therapeutic target that could be leveraged alongside strategies aimed at regulating *SOX10*.

Most significantly, our findings also reveal specific enhancer elements essential for *SOX10* regulation and phenotype switching in melanoma. By manipulating *SOX10* expression and identifying the role of NTRK1 in drug-resistant phenotypes, we propose a dual-targeting strategy that could disrupt melanoma’s adaptive capacity, potentially eradicating cells that survive conventional therapies. This work not only advances our understanding of melanoma biology but also opens avenues for targeted therapies against metastatic melanoma, where interventions targeting *SOX10* regulatory pathways, MAPK, and NTRK1 could provide a much-needed therapeutic advantage.

## Results

### Loss of a specific sox10 enhancer alters melanoma onset rate in a zebrafish model

Recognizing *SOX10*’s key role in NC and melanoma cell identity, we sought to elucidate the transcriptional mechanisms controlling its expression and investigate how melanoma reactivates the embryonic gene program in the context of oncogenesis^3^. We used the well-established BRAF^V600E^;p53^lof/lof^ zebrafish melanoma model, in which the most common human BRAF oncogene coding sequence (V600E) is expressed in a melanocyte-specific manner under the control of the zebrafish *mitfa* promoter with a global p53 loss-of-function mutation^41^. These genetically engineered zebrafish models all develop melanomas with histologic and molecule features highly analogous to human melanoma. In our prior work, we identified chromatin regions with differential accessibility by ATAC-seq in zebrafish melanoma tumor cells compared to melanocytes, with each differentially accessible peak representing a putative enhancer element reactivated in melanoma^40^.

We have previously shown that specific evolutionarily-conserved enhancers upstream of *sox10*, in particular *peak5* (a 669 bp region 15 kb upstream of the *sox10* transcriptional start site), are required for wild type levels of *sox10* expression during embryonic development and for precise melanocyte patterning in zebrafish^40^ (**Supp Fig. 1a, b**). However, these zebrafish develop and breed otherwise normally. Given the central importance of *sox10* expression in zebrafish and human melanoma formation and growth^42–44^ and the melanoma-specific reporter activity of *peak5*- driven EGFP reporters, we wondered if deletion of *peak5* would also alter the rate of *de novo* melanoma onset in our *BRAF-*driven melanoma zebrafish model^41^. We bred the homozygous deletion of *peak5* (*stl538*) allele (**Supp Fig. 1c**) that we previously generated into the BRAF/p53 zebrafish melanoma model^3,41^, and found this significantly delayed melanoma onset (median 297.5 days, homozygous *peak5* deletion) as compared to heterozygous deletion of *peak5* or wild type (median 273 days) (Gehan-Breslow-Wilcoxon test P value of 0.0447 Mantel-Hänszel Hazard Ratio of 1.609) **(Fig. 1a)**. These results indicate that enhancers for *sox10*, like *peak5*, can be deleted or their inputs potentially inhibited, and remain compatible with largely normal development while also having important melanoma-specific activity in regulating tumor onset. This further supports the utility of analyzing individual enhancer elements for their potential specific roles in regulating *sox10* levels/activity in different contexts.

**Figure 1.**
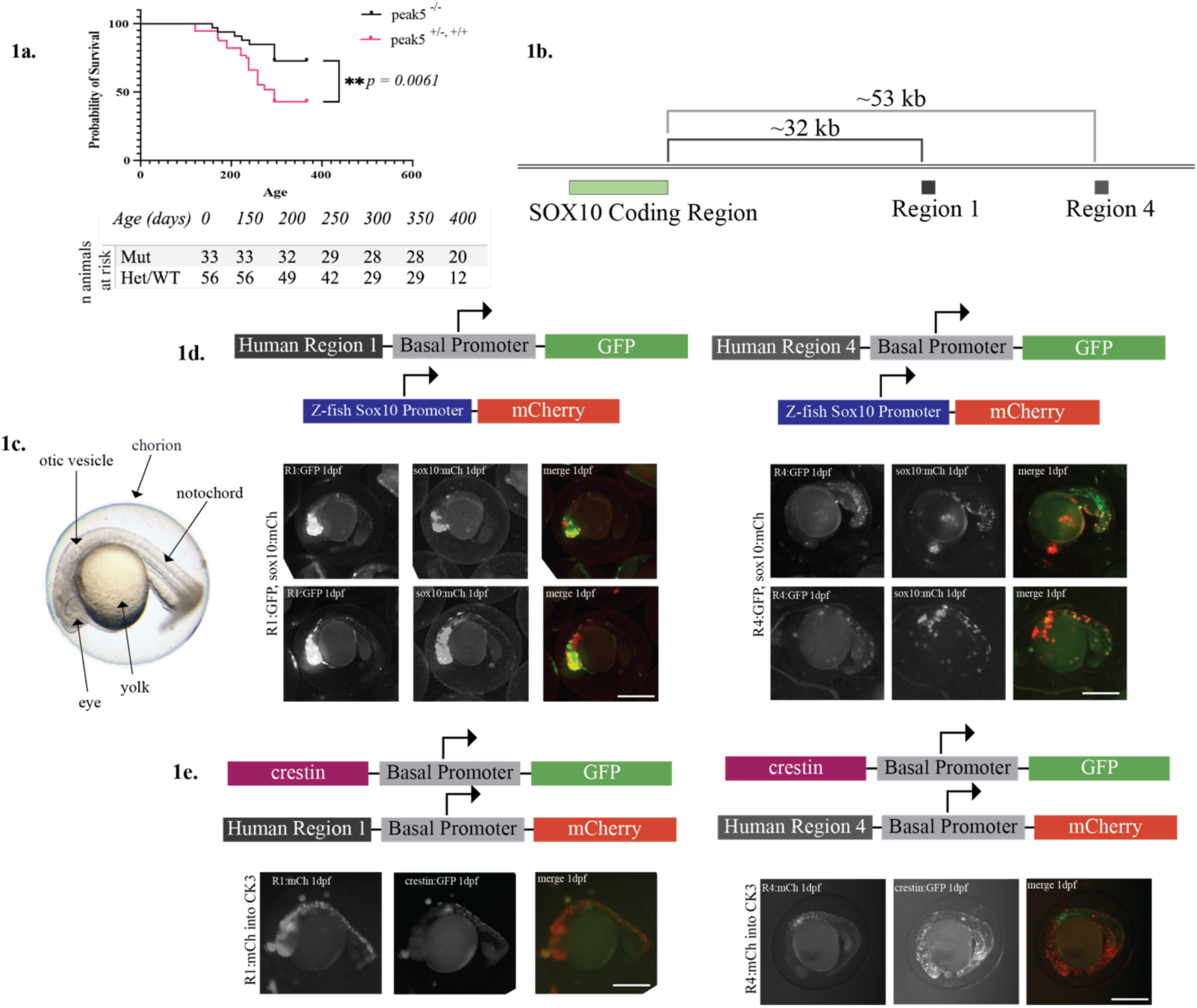
Identification and Functional Testing of Melanoma-Associated Enhancers in Zebrafish 1a. Kaplan-Meier survival analysis of BRAF-driven zebrafish melanoma models, comparing survival probabilities between wild-type and peak5-deleted genotypes. A significant delay in melanoma onset is observed in the homozygous peak5 deletion group (median survival: 297.5 days) compared to wild-type and heterozygous deletion (median survival 273 days). **1b.** Schematic representation of key human SOX10 regulatory regions selected from cross-species analysis. The SOX10 coding region is located approximately 32 kb upstream of Region 1, and approximately 53 kb upstream of Region 4). **1c.** Bright field image of 1 dpf zebrafish embryo with key anatomical features annotated. **1d.** Transgenic zebrafish embryos expressing GFP and mCherry reporter constructs driven by human SOX10 enhancer regions. (Left): shows expression of the GFP reporter for human Region 1 and the zebrafish sox10 promoter-driven mCherry at 1dpf. (Right): shows expression of the GFP reporter for human Region 4 and the zebrafish sox10 promoter-driven mCherry at 1dpf. **1e.** Transgenic zebrafish embryos expressing GFP and mCherry reporter constructs driven by human SOX10 enhancer regions. (Left): shows expression of mCherry driven by human enhancer Region 1 with overlapping expression of crestin:GFP at 1dpf. (Right): shows expression of mCherry driven by human enhancer Region 4 with overlapping expression of crestin:GFP at 1dpf.

### Translating zebrafish to human regulatory regions

Given that the majority of zebrafish enhancer elements, including those regulating *sox10*, do not exhibit extended regions of linear sequence conservation with human *SOX10* regulatory elements, we sought to identify alternative defining features of key zebrafish enhancers that could be extrapolated to human contexts.

Despite the absence of extended stretches of sequence conservation between zebrafish and human, we adapted our zebrafish analytical framework for human enhancer identification by first identifying differentially accessible regulatory regions in melanoma. In the zebrafish, our search for melanoma-specific enhancer elements began with regions of differential chromatin accessibility, as determined by ATAC-seq (**Supp Fig. 2a, left**). In the human genome, we initially identified regions of potential regulatory interest based on specific H3K27ac marks^6^, a well-established indicator of active enhancers, across multiple melanoma cell lines (**Supp Fig. 2a, right**).

**Figure 2.**
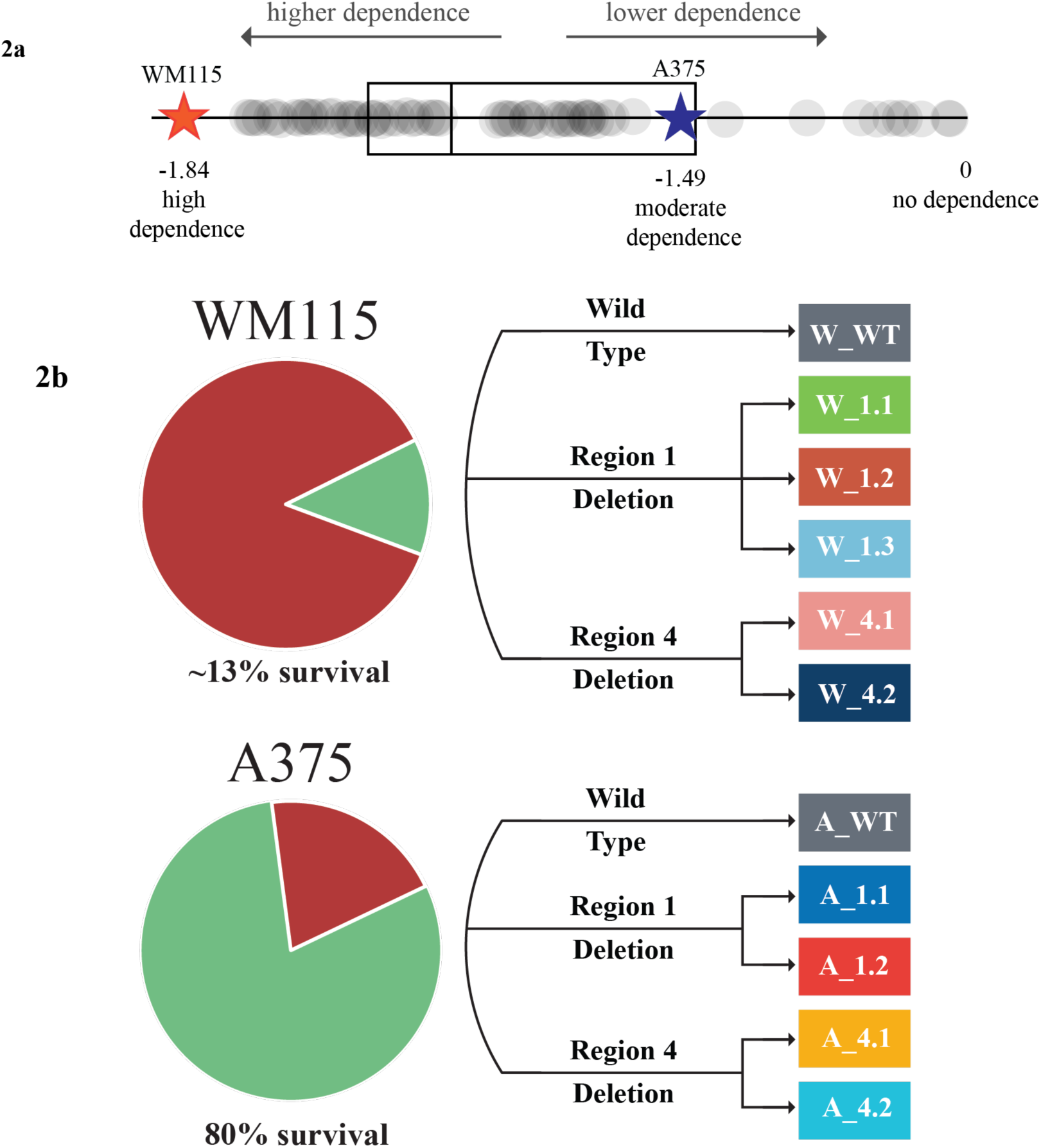
SOX10 Dependency and Enhancer Deletion in Melanoma Cell Lines 2a. Schematic illustrating SOX10 dependency in human melanoma cell lines based on Chronos dependency scores. WM115 cells exhibit a high dependency on SOX10, while A375 cells show only moderate dependency. **2b.** Generation of enhancer deletion lines. CRISPR/Cas9 and guide RNAs targeting enhancer elements were electroporated into both cell lines, followed by single cell sorting and genotyping to confirm successful deletion. Tolerance of WM115 and A375 of the targeted deletion of enhancer elements. **2b (top):** The highly SOX10 dependent WM115 cells showed low tolerance for enhancer deletion, with only ∼13% of clones harboring the deletion surviving. **2b (bottom):** The less SOX10-dependent A375 cells demonstrated higher tolerance to enhancer deletion, with ∼80% of the deletion clones surviving. 11 stable deletion lines were generated: each line is labeled as follows: ‘Parental Cell Line’_‘Region Number’. ‘Replicate Number’. Additionally, wild type (WT) A375 and WM115 cells underwent the same process of single-cell sorting and clonal expansion to serve as controls, ensuring consistency in selection conditions.

As in the zebrafish analysis, we refined the candidate regions by evaluating evolutionary conservation for more closely related vertebrate species as conserved sequences often represent regions of functional significance. By aligning the zebrafish sox10 regulatory region with those of related Cyprinidae (carp) species, we identified only a handful of conserved sequences, as visualized by the dot plot (**Supp Fig. 2b, left**). These conserved sequences overlapped with a subset of peaks identified by the ATAC-seq, as marked (Peak 2/3, Peak4, Peak5, Peak8)^40^. For the human analysis, we applied a similar approach by conducting comparative genomic analyses against multiple vertebrates (chicken, rat, mouse, ape) and identified four regions with conserved sequence elements (**See Supplementary Table 1**). The dot plot in **Supp Fig. 2b, right,** illustrates the alignment between the human *SOX10* locus and the corresponding rat region.

Given our previous findings that closely spaced SOXE binding motifs (two binding sites separated by 3-5 nucleotides) are crucial for the neural crest and melanoma-specific activity of the zebrafish peak5^40^ (**Supp Fig. 2c, left**), we screened the four conserved human regions for similar SOXE dimer sites. Only two evolutionarily conserved regions within the 60 kb putative upstream regulatory region of human *SOX10* satisfied all criteria, referred to henceforth as “Region 1”; (501 bp region 31,947bp upstream of the sox10 coding start site) and “Region 4” (500 bp region 53,088 upstream of the sox10 coding site) (**Fig. 1b)**. Upon comparison with existing datasets, we found that an MPRA study^45^ highlighted these regions as putative enhancers and SOX10 ChIP-seq^46^ confirmed the presence of *SOX10* dimer sites at these coordinates. Consequently, we selected these two regions for further investigation.

As the presence of key regulatory transcription factor binding sites (TFBS) has been shown to drive conserved functions across species^47,48^, we wondered if these human enhancer elements (Region 1 and Region 4) would drive spatially and temporally similar reporter expression as zebrafish *sox10* transcriptional control elements and other neural crest markers/reporters (e.g. the well-characterized zebrafish neural marker *crestin*)^3^, despite the lack of apparent extended sequence conservation. We generated multiple independent transgenic F0 reporter zebrafish embyos using standard Tol2-based methods **(Fig. 1c)** (which efficiently yields random insertion transgenic animals) by coinjecting reporters for human *SOX10* enhancers [*Region1:EGFP* **(Fig. 1c, left)** or *Region4:EGFP* **(Fig. 1c, right**)] and zebrafish *sox10* enhancers (*sox10_MP:mCh.*) Additionally, we injected reporters for human *SOX10* enhancers (*region1:EGFP* **(Fig. 1d, left)** or *region4:EGFP* **(Fig. 1d, right**) into a line bearing stable expression of a neural crest reporter construct (*crestin:mCh*)^3^. Remarkably, we found significant overlap of reporter expression for both human-and zebrafish-specific neural crest reporters with the human enhancer elements target EGFP to neural crest cells **(Fig. 1c and 1d**). This further supports the hypothesis that these human enhancer elements may represent functional developmental *Sox10* regulatory elements that drive neural crest-specific gene expression and play important roles in human melanoma transcriptional programs.

### Deletion of key human Sox10 enhancer elements and effects on melanoma growth

As loss of a key enhancer element of *sox10* in the zebrafish model caused a significant delay in melanoma onset (**Fig 1a)**, we sought to explore whether targeted deletion of enhancer elements sharing key characteristics (i.e. presence of a conserved SOXE dimer, vertebrate sequence conservation as found for Regions 1 and 4, and NC-specific reporter activity in developing zebrafish) in human melanoma cell lines would have similar anti-melanoma effects. Using the depmap resource, we selected melanoma cell lines with a range of *SOX10* dependencies including A375 cells (Chronos Gene Effect score of-1.49) and WM115 cells (Chronos Gene Effect score of-1.84, where a score of 0 indicates no dependence and a more negative score indicates a higher dependency)^49^ **(Fig. 2a)**.

A375 and WM115 cells were electroporated to introduce CRISPR/Cas9 and gRNAs targeting either Region 1 or Region 4 **(Fig. 2b)**. Single cells were sorted, and genetically altered populations were grown from a single clone. The WT counterparts were also put through this single cell selection bottleneck. In A375 cells, targeted deletion of either enhancer element via CRISPR/Cas9-mediated deletion was moderately well-tolerated, with approximately 80% of clones demonstrating stable growth post-deletion (**Fig. 2b, top)**. In contrast, the more highly *SOX10*-dependent WM115 cells exhibited a much lower tolerance for enhancer loss, with only 13% of clones able to survive the deletion of a *Sox10* enhancer. **(Fig. 2b, bottom)**.

We generated multiple stable deletion lines for each targeted region in each cell line, using a consistent naming convention for clarity. Each line is labeled as follows: *‘Parental Cell Line’_‘Region Number’. ‘Replicate Number’.* For example, “A_4.1” refers to the first stable deletion line for Region 4 in the A375 cell line. For each targeted region, we generated two to three stable lines per parental cell lines **(Fig. 2b)**.

We first showed that deletion of Region 1 or Region 4 led to significant downregulation of *SOX10* expression as measured by qPCR in almost all the deletion lines (Welch’s ANOVA test P value of 0.0005 in WM115 lines, Welch’s ANOVA test P value of <0.0001 in A375 lines) (**Fig. 3a, 3b; left)**. These changes were also reflected in protein levels as western blots for SOX10 also showed significant reduction in protein, notably in line W_1.2 (**Supp. Fig. 3)**. These data indicate the importance of these enhancer regions for *SOX10* transcriptional activity and establishes their identification as *bona fide* enhancers of *SOX10* endogenous expression in human melanoma.

**Figure 3.**
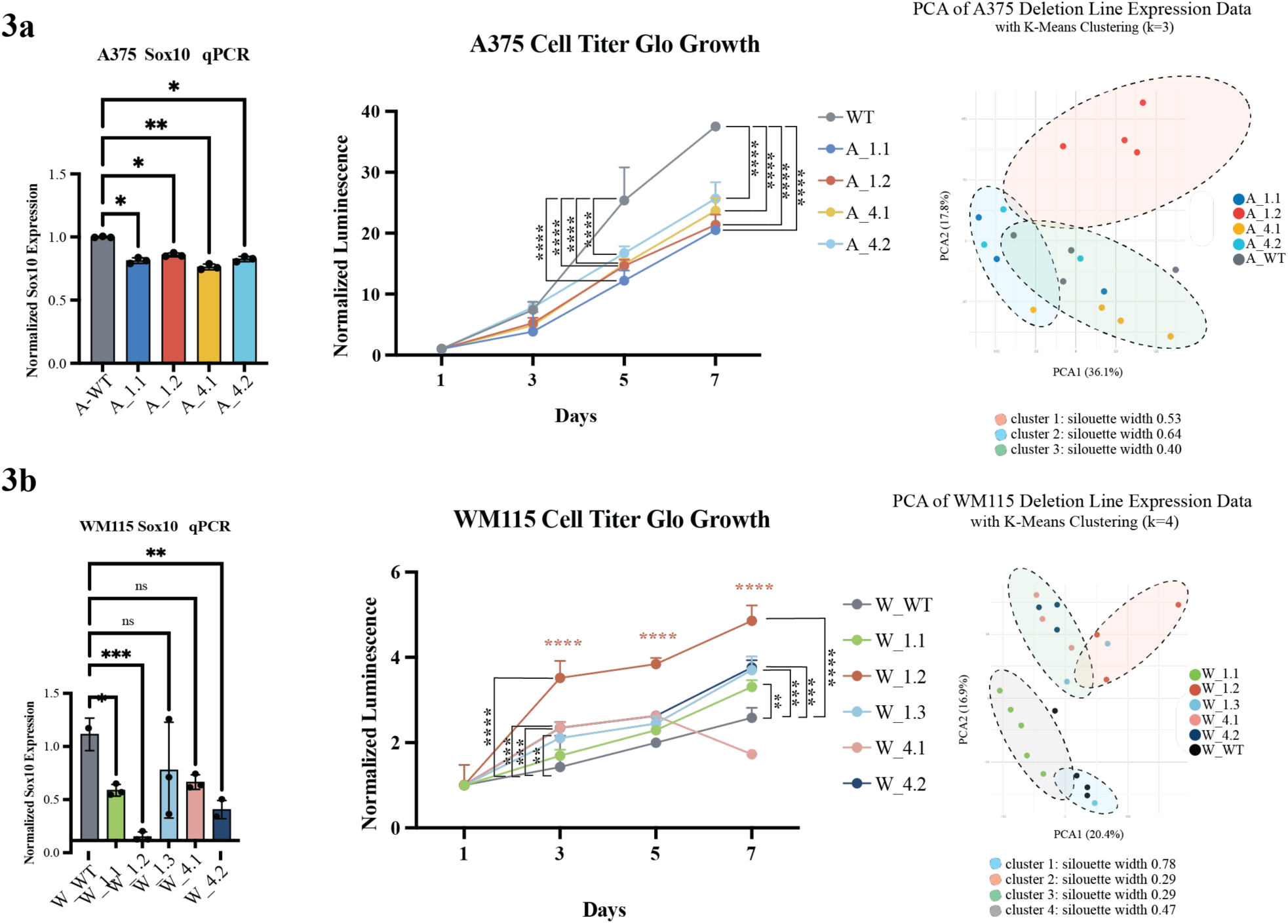
Effects of Enhancer Deletion on SOX10 Expression and Cell Proliferation in A375 and WM115 Cell Lines 3a. A375 Cell Lines (Left) Normalized SOX10 mRNA expression levels measured by qPCR in A375 cells with and without SOX10 enhancer deletions (A_WT, A_1.1, A_1.2, A_4.1, A_4.2). Deletion of enhancer elements resulted in significant reductions in SOX10 expression in several lines, with varying magnitudes. Statistical significance: *p < 0.05, **p < 0.01. **(Center)** Proliferation curves of A375 cell lines measured by CellTiter-Glo luminescence assay over 7 days. Deletion lines showed slightly reduced proliferation compared to the A_WT control. Statistical significance across time points: ****p < 0.0001. **(Right)** Principal Component Analysis (PCA) of A375 deletion line RNA-seq data, colored by k-means clustering (k=3). Clustering reflects subtle shifts in gene expression profiles across deletion lines, silhouette width cluster optimization. **3b. WM115 Cell Lines (Left)** Normalized SOX10 mRNA expression levels measured by qPCR in WM115 cells with and without SOX10 enhancer deletions. Deletion of enhancer elements significantly reduced SOX10 expression in most lines. Statistical significance: *p < 0.05, **p < 0.01, ***p < 0.001, ****p < 0.0001, ns: not significant. **(Center)** Proliferation curves of WM115 cell lines measured by CellTiter-Glo luminescence assay over 7 days. Deletion lines exhibited significantly increased proliferation relative to W_WT. Statistical significance across time points: ****p < 0.0001. **(Right)** PCA of WM115 deletion line RNA-seq data, colored by k-means clustering (k=4). Clustering highlights distinct shifts in transcriptional profiles, reflecting differentiation state transitions upon enhancer deletion, silhouette width cluster optimization.

Knockdown of *SOX10* mRNA in melanoma cells has been shown to induce senescence, cell death, and reduced growth rates^50–52^. We therefore hypothesized that deletion of these enhancer elements and consequent decreased *SOX10* expression (**Fig. 3a,b)** would similarly impact cell growth. In A375 Region1 and Region 4 deletion lines, we measured cellular proliferation (CellTiter Glo) and found that deletion lines exhibited significantly slower growth rates compared to their WT counterpart consistent with this hypothesis, with growth rate fold changes ranging from a 0.23-fold decrease (A_1.2) to a 0.13-fold decrease (A_4.2) at day 7 compared to their WT counterparts (all lines shown in **Fig. 3a, middle**).

Interestingly, deletion of Regions 1 and 4 in WM115 melanoma cells, which depmap predicted to have high *SOX10* dependency, led to an unexpected increase in growth rate in deletion lines, despite significantly reduced *SOX10* expression (**Fig. 3b, left**). WM115 deletion lines showed generally faster growth rates than their WT counterparts, with increases ranging from a 1.27-fold change (W_4.1) to a 3.53-fold change (W_1.2) at day 7 compared to WT (**Fig. 3b, middle**). Indeed, proliferation rates correlated negatively with *SOX10* expression levels in WM115 cells (correlation coefficient-0.852, R^2^=0.73), indicating that gene expression programs tolerant of decreased *SOX10* expression paradoxically facilitated faster growth (**Supp. Table 2**). This, together with the lower clonability observed in the WM115 cells during the CRISPR engineering process, suggests that these highly *SOX10-*dependent cells adapted to the pressure of reduced *SOX10* expression, possibly by engaging alternative pathways to support enhanced proliferation. In contrast, A375 cells, with only moderate *SOX10* dependency, showed the expected moderate reduction in growth rate, potentially requiring fewer adaptive changes in response to decreased *SOX10* expression.

### Transcriptional adaptations following deletion of Region 1 and 4 SOX10 enhancers

To explore how melanoma cells transcriptionally adapt to the engineered *SOX10* enhancer deletions, we analyzed the transcriptomes from each independent line using bulk RNA-seq. As described above (**Fig 2b**), 11 lines were generated from A375 and WM115 cells. Total RNA was extracted from each and sequenced in bulk, with 3 to 6 biological replicates per line. When comparing all deletion lines to their WT counterparts, we observed a consistent transcriptomic shift, as visualized in the PCA plots. The PCA of A375 deletion lines formed three distinct clusters (k = 3, average silhouette width = 0.51), with lines like A_1.2 clustering separately from their A_WT counterparts (**Fig. 3a, right**). The WM115 deletion lines grouped into four main clusters (k = 4, average silhouette width = 0.47), with the W_WT line clustering near the W_1.1 cluster, and the remaining deletion lines clustering together separately (**Fig. 3b, right**).

In addition, we noted significant alterations in the bulk RNA-seq analysis of genes known to be related to melanoma phenotype switching including SOX10, SOX9, MITF, and AXL, among others^18,53,54^. These genes have been linked to phenotypic alterations of proliferation and migratory ability as defined in Hoek et al and Rambow et al^55,56^. We thus examined both WM115 and A375 deletion lines relative to their WT counterparts for changes in gene signatures related to proliferative/melanocytic and invasive/mesenchymal phenotypes. Indeed, deletion of either region in both WM115 and A375 lines displayed a downregulation of genes (such as SOX10, MITF, and ZEB2) that have been implicated in maintaining the proliferative/melanocytic phenotype and a consequent upregulation of the phenotype switch related genes (such as SOX9, AXL, and ZEB1) supporting a more invasive/mesenchymal phenotype^54^ (**Supp.** Fig. 4).

To assess phenotype shifts in each deletion line in a quantitative manner, we scored the expression of gene lists defining Tsoi sub-phenotype categories, generating a weighted trajectory position score ranging from 1 (melanocytic) to 7 (undifferentiated)^19^. These trajectory scores, summarized in **Table 1** and **Figure 4a**, provide a snapshot of one way of defining phenotype scores, as previously published by Tsoi et al. in 2018. While we acknowledge that phenotypes likely encompass factors beyond these gene lists, the scores reveal a clear trend across all deletion lines: the loss of SOX10 enhancer elements shifts the phenotype identity toward a more undifferentiated state. Interestingly, the baseline WM115 line (W_WT) scored lower on the trajectory (3.569, SD = 0.251) than the baseline A375 line (A_WT) at 4.431 (SD = 0.146), as the higher SOX10 dependence in the WM115 cells supports a more melanocytic phenotype. While not all lines reached statistical significance in their mean score difference or score shift, 8 out of the 9 deletion lines exhibited a shift towards a more undifferentiated phenotype score (**Fig 4b, see Supplementary Table 3 for detailed information)**. A heatmap of individual category scores across deletion lines is shown in **Fig. 4c**.

**Figure 4.**
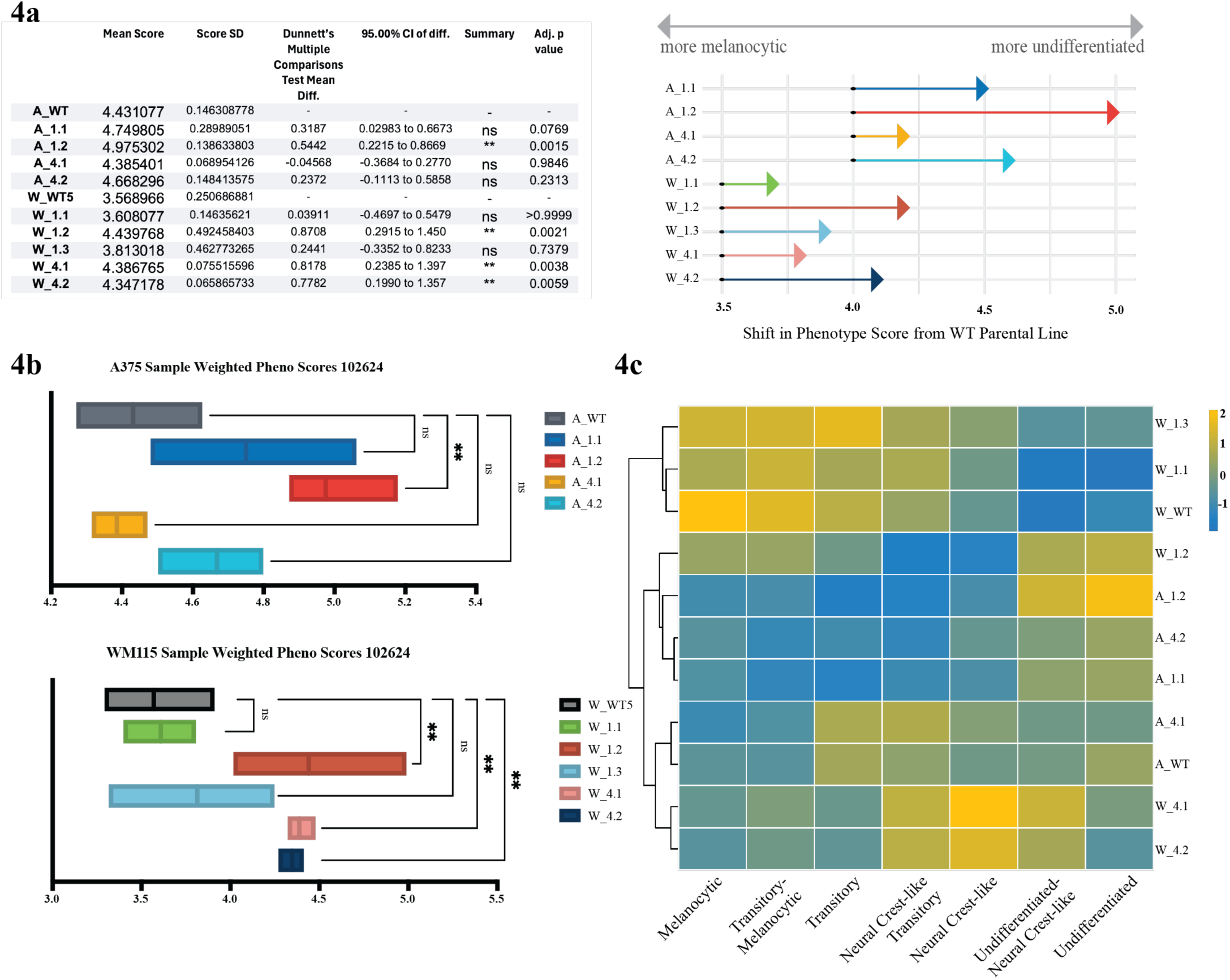
Phenotype scores and expression profiles highlight differential trajectories and gene expression across deletion lines. Fig 4a. Phenotype trajectory scores across deletion lines: Mean phenotype trajectory scores and statistical comparisons (Dunnett’s test) for A375 and WM115 cell lines with and without SOX10 enhancer deletions, based on weighted expression of Tsoi sub-phenotype gene lists (melanocytic to undifferentiated, 1–7). Scores for WM115 deletion lines (W_1.2, W_4.1, W_4.2) show significant shifts towards more undifferentiated phenotypes compared to W_WT. In A375, only A_1.2 exhibits a statistically significant shift, though trends are evident in A_4.1 and A_4.2. Arrow plot **(right)** visualizes the magnitude and direction of shifts in phenotype scores relative to WT parental lines. Fig 4b. Boxplots of weighted phenotype scores: Weighted phenotype trajectory scores for A375 (top) and WM115 (bottom) deletion lines. WM115 lines exhibit more pronounced and statistically significant shifts towards undifferentiated states compared to A375 lines. Adjusted p-values: **p < 0.01, ***p < 0.001, ns: not significant. Fig 4c. Heatmap of phenotype category scores: Heatmap of individual sub-phenotype category scores (adapted from Tsoi et al. gene lists) for each deletion line.

### Response to BRAF/MEK inhibitor therapy

Since phenotype switching has been linked to responsiveness to BRAF and MEK inhibitors, we next investigated whether the loss of specific *SOX10* enhancer elements and the resulting phenotype shift could contribute to drug resistance in human melanoma cells^57–59^. We treated each A375 and WM115 Region 1 and Region 4 deletion line with dabrafenib (a BRAF inhibitor) or trametinib (a MEK inhibitor) across a concentration range of 0.001-1000 nM to determine IC50 values. Deletion lines derived from both the A375 and WM115 parental cell lines have increased IC50 values, indicating a greater capacity to tolerate these common melanoma treatments (see **Fig 5** and **Supplementary Tables 4 and 5** for specific data). For instance, the IC50 for dabrafenib increased dramatically from 0.5594 in W_WT cells to 78.24 in W_1.2 cells (**Fig 5b, left)**. Similarly, in response to trametinib, IC50 values increased from 0.1454 in W_WT cells to 6.718 in W_1.2 cells (**Fig 5b, right)**.

**Figure 5.**
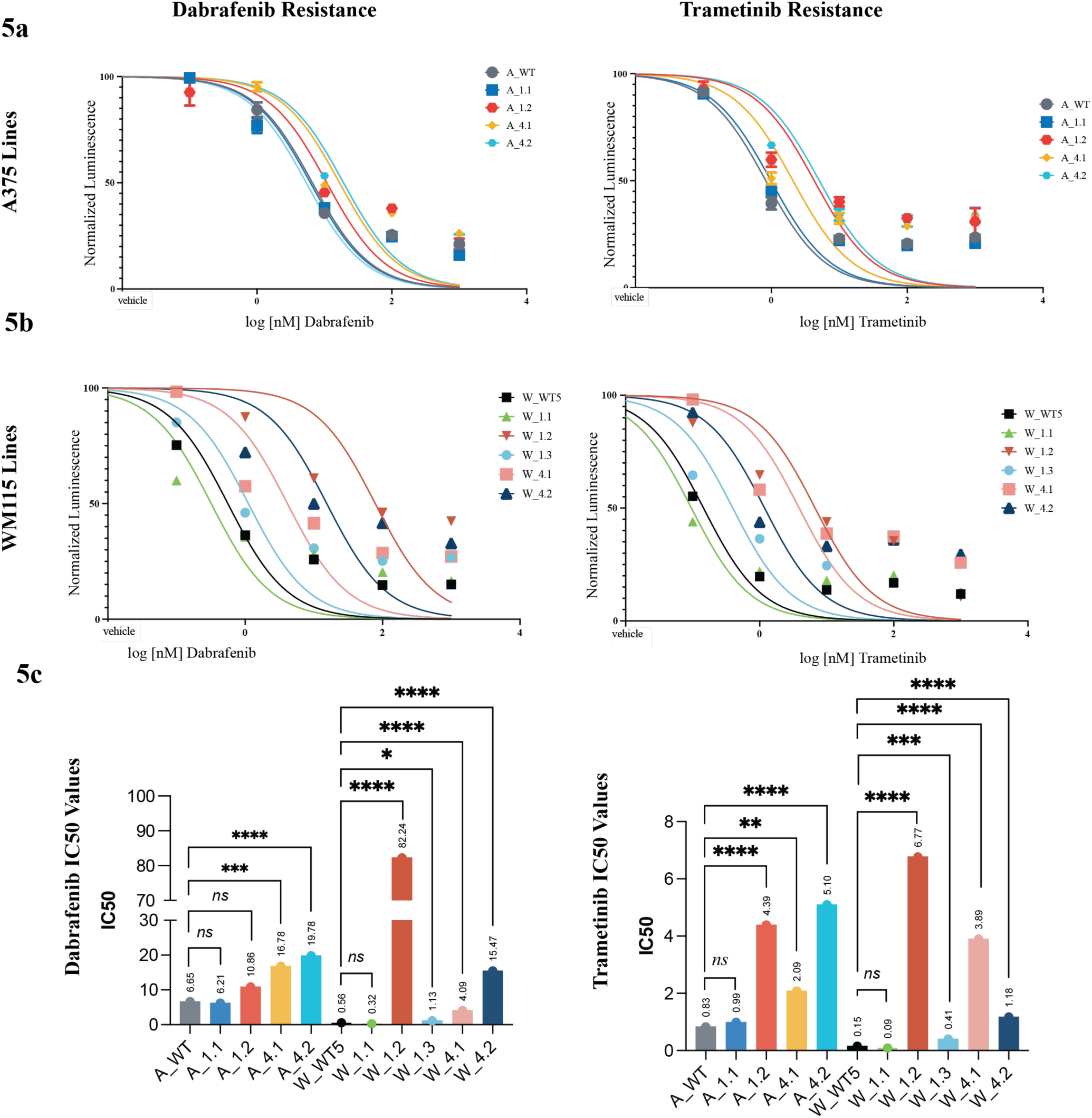
Effect of Enhancer Deletion on Drug Resistance (5a) Cell viability of A375 cells challenged with (left): dabrafenib and (right): trametinib. All lines show an increased IC50, indicating greater drug resistance compared to their WT counterparts. **(5b)** Cell viability of WM115 cells challenged with (left): dabrafenib and (right): trametinib. Similar to A375, almost all deletion lines exhibit increased IC50 values, suggesting a higher level of drug resistance than the WT. **(5c)** Bar graphs showing the IC50 values for each line under both drug conditions. IC50 values for each deletion line are plotted for dabrafenib (left), and trametinib (right), highlighting the differences in drug resistance across the various lines.

Deletion lines that underwent the least substantial shifts in phenotype based on changes in gene expression as assessed above (**Fig 4)**, such as A_4.1 and W_1.1, consistently displayed the lowest IC50 values (highest drug sensitivity) for both drugs, responding similarly to their WT counterparts. In contrast, lines that shifted most significantly towards an undifferentiated fate (A_1.2, W_4.1, and W_4.2) showed the highest IC50 values (highest levels of resistance) to both dabrafenib and trametinib (**Fig 5c)**. A slight position correlation was observed between the numeric trajectory position score and resistance to dabrafenib (R^2^ = 0.67 sans W_1.2, R^2^ = 0.293 including the W_1.2 extreme value) and trametinib (R^2^ = 0.7). When looking at the mean difference of score, or the amount that the score shifted, the correlation was slightly more with dabrafenib (R^2^ = 0.589 sans W_1.2, R^2^ = 0.44 including the W_1.2 extreme value) and trametinib (R^2^ = 0.614) (**Supplementary Table 6)**. These data indicate that deletion lines exhibiting minimal phenotype shifts away from a melanocytic state tend to maintain drug sensitivity, while those shifting towards an undifferentiated state develop increased resistance.

### Modulation of SOX10 levels and targeted drug resistance pathways

To more broadly assess associated gene regulatory changes associated with modulating *SOX10* activity via deletion of specific enhancer elements, we performed GSEA analysis of 7 human gene set collections encompassing 29,316 gene sets from GSEA-misgdb individually for each deletion line, then analyzed for shared up and down regulated pathways across lines (see methods). The highest represented pathways in all deletion lines included gene sets shown to be important for melanoma metastasis^60^, melanoma relapse^61^, cell migration^62^, and EMT^63^. **(Supp.** Fig. 5). Overall, these data again link alterations of SOX10 levels via enhancer deletions from a more melanocytic phenotype to a shift towards an invasive/mesenchymal phenotype.

To investigate the mechanisms underlying drug resistance, we focused on differentially expressed genes associated with the *SOX10* enhancer deletions we generated, which may play a key role in producing the observed BRAF/MEK inhibitor resistance phenotype. Across all deletion lines, an average of 1,800 genes were significantly upregulated (Supp Figure 5B), with cell lines showing a larger phenotype score shift exhibiting a higher number of upregulated genes (Pearson correlation score 0.748), suggesting that transcriptional reprogramming drives this phenotype shift.

We prioritized a subset of these genes based on their differential expression and relevance to MAPK inhibitor (MAPKi)-resistant melanoma tumors isolated from patients^64^ or the presence of an invasive phenotype motif across cell lines^18^ (Supp Fig 6). Genes were ranked using a weighted priority score calculated from a combination of log fold change, adjusted p-value thresholds, and observed IC50 values for dabrafenib and trametinib. AMIGO2 (score: 17.9), RCOR2^65,66^ (score: 17.6), NTRK1 (score: 17.4), and NKX3.1 (score: 16.8) were among the top 40 prioritized genes (**Fig. 6a**), showing strong potential relevance to melanoma resistance mechanisms and phenotype switching.

**Figure 6.**
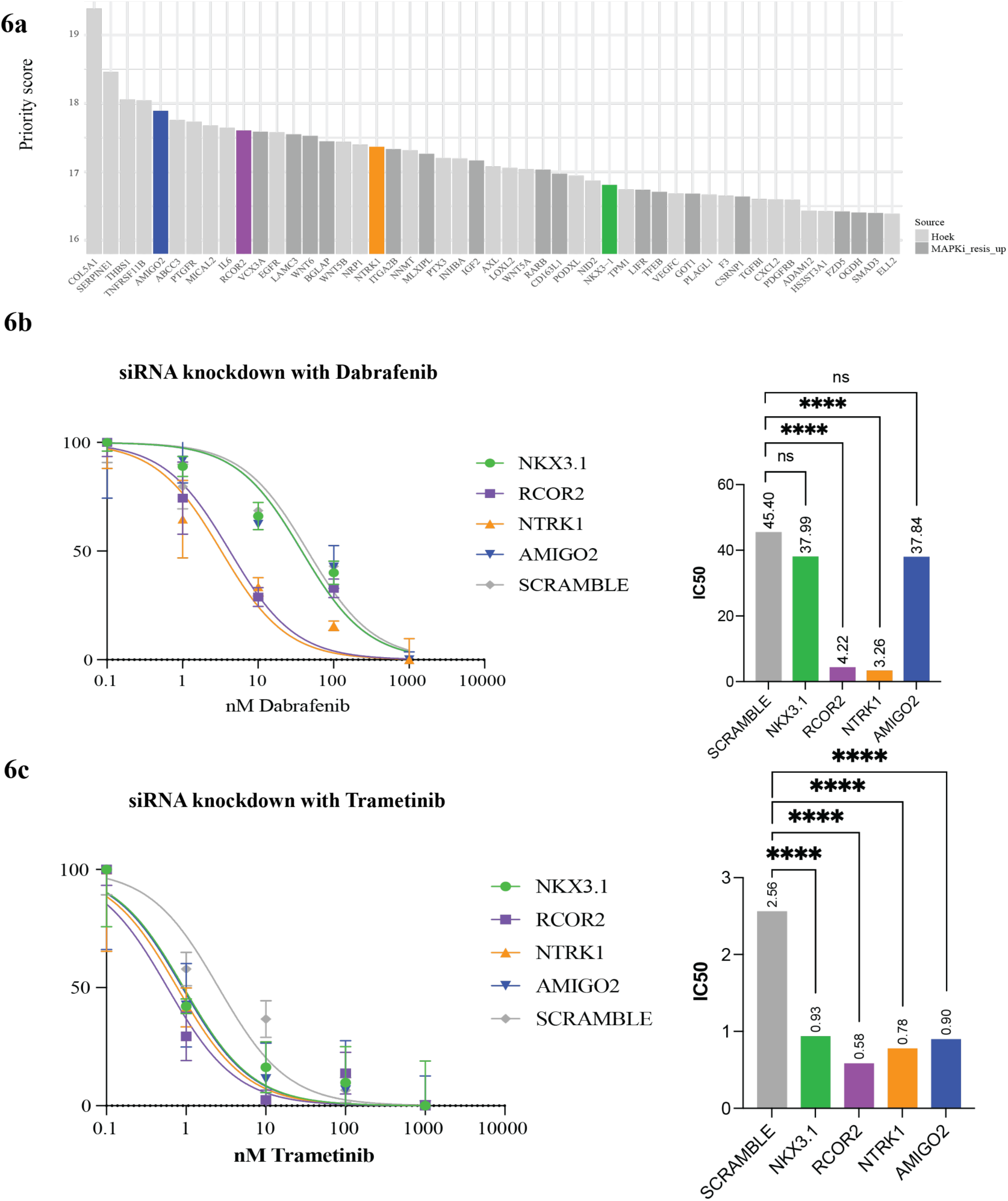
Target gene identification and validation in MAPKi resistance 6a. Bar plot displaying the priority scores for the top 50 genes ranked from genes upregulated in MAPKi resistant melanoma or Hoek invasive motif gene sets. Key genes of interest (NKX3.1, RCOR2, NTRK1, AMIGO2) are highlighted in distinct colors. **6b.** IC50 curves and bar plot for dabrafenib sensitivity. **(Left):** Dose-response curves for dabrafenib in cell lines with knockdown of NKX3.1, RCOR2, NTRK1, AMIGO2, and a scramble control. IC50 values were calculated based on viability assays, revealing increased resistance for RCOR2 and AMIGO2 knockdowns compared to scramble. **(Right):** Bar plot of IC50 values (nM) for dabrafenib treatment across the knockdown lines. NTRK1 and RCOR2 knockdown significantly reduced IC50 values. **6c.** IC50 curves and bar plot for trametinib sensitivity. **(Left):** Dose-response curves for trametinib in cell lines with knockdown of NKX3.1, RCOR2, NTRK1, AMIGO2, and a scramble control. IC50 values demonstrate differential sensitivity, with RCOR2 knockdown exhibiting the highest resistance. **(Right):** Bar plot of IC50 values (nM) for trametinib treatment across knockdown lines. All siRNA restored trametinib sensitivity significantly compared to the scramble control.

phenocopy their impact on drug sensitivity in a representative cell line. Knockdown of RCOR2 and NTRK1 yielded increased sensitivity to dabrafenib, restoring dabrafenib IC50 from 10.86 nM in the A_1.2 line to 4.22 nM and 3.26 nM, respectively. Notably, this level of sensitivity was even greater than the IC50 for dabrafenib in the A_WT line of 6.65 nM (**Fig. 6b**). Interestingly, NTRK1 was upregulated in nearly all other phenotype-switched lines, and while expressed in line A_1.2, was slightly downregulated (-0.644 logFC), overall suggesting that NTRK1 is part of a broader gene regulatory network influencing drug resistance, potentially independent of its own baseline expression in melanoma cells.

Additionally, when these genes were knocked down in trametinib resistant A_1.2 cells (IC50 = 4.39), NTRK1 knockdown was the only gene to restore the trametinib sensitivity to below that of the A_WT line (IC50 = 0.78 compared to IC50 = 0.83), highlighting NTRK1’s potential contribution to drug resistance in melanoma (**Fig 6b)**.

To further explore the role of NTRK1 in drug resistance, we examined clinical data from BRAF/MEK inhibitor resistant melanoma samples, where NTRK1 upregulation was frequently observed. Our analysis showed that NTRK1 expression correlated more strongly with phenotype scores (R˄2 = 0.714) than *SOX10* expression (R˄2 =-0.09) (**Supplementary Table 6)**. When we referenced genes upregulated in MAPKi-exposed melanoma tumors, NTRK1 was significantly upregulated in 7 out of 9 deletion lines. These findings underscore NTRK1’s potential involvement in melanoma phenotype switching and resistance to targeted therapies.

Next, we tested whether NTRK1 inhibition could impact drug resistance in the A_1.2 line, a resistant, phenotype-shifted line that showed high resistance to both dabrafenib and trametinib despite lacking top-level NTRK1 upregulation. Treatment with NTRK inhibitors entrectinib^67,68^ and larotrectinib^69^ in combination with BRAF/MEK inhibitors resulted in an additive effect, reducing the doses needed to achieve cell death, with certain combinations of dabrafenib and entrectinib achieving ZIP scores above 10 indicating synergy (**Fig 7a)**. Dose-response matrices and ZIP synergy score plots for A_WT and A_1.2 cell lines challenged with combinations of dabrafenib (BRAFi) and entrectonib (NTRKi). Dose-response matrices (**Fig. 7b, left**) show the percentage inhibition across a gradient of dabrafenib and entrectonib doses. ZIP synergy score plots (**Fig. 7b, right**) demonstrate areas of synergy (positive scores). A_WT exhibits minimal synergy with the combination treatment (ZIP synergy score median =-3.18), whereas A_1.2 shows pronounced synergy, with a maximum ZIP synergy score of 13.68 (ZIP synergy score median = 1.52).

**Figure 7.**
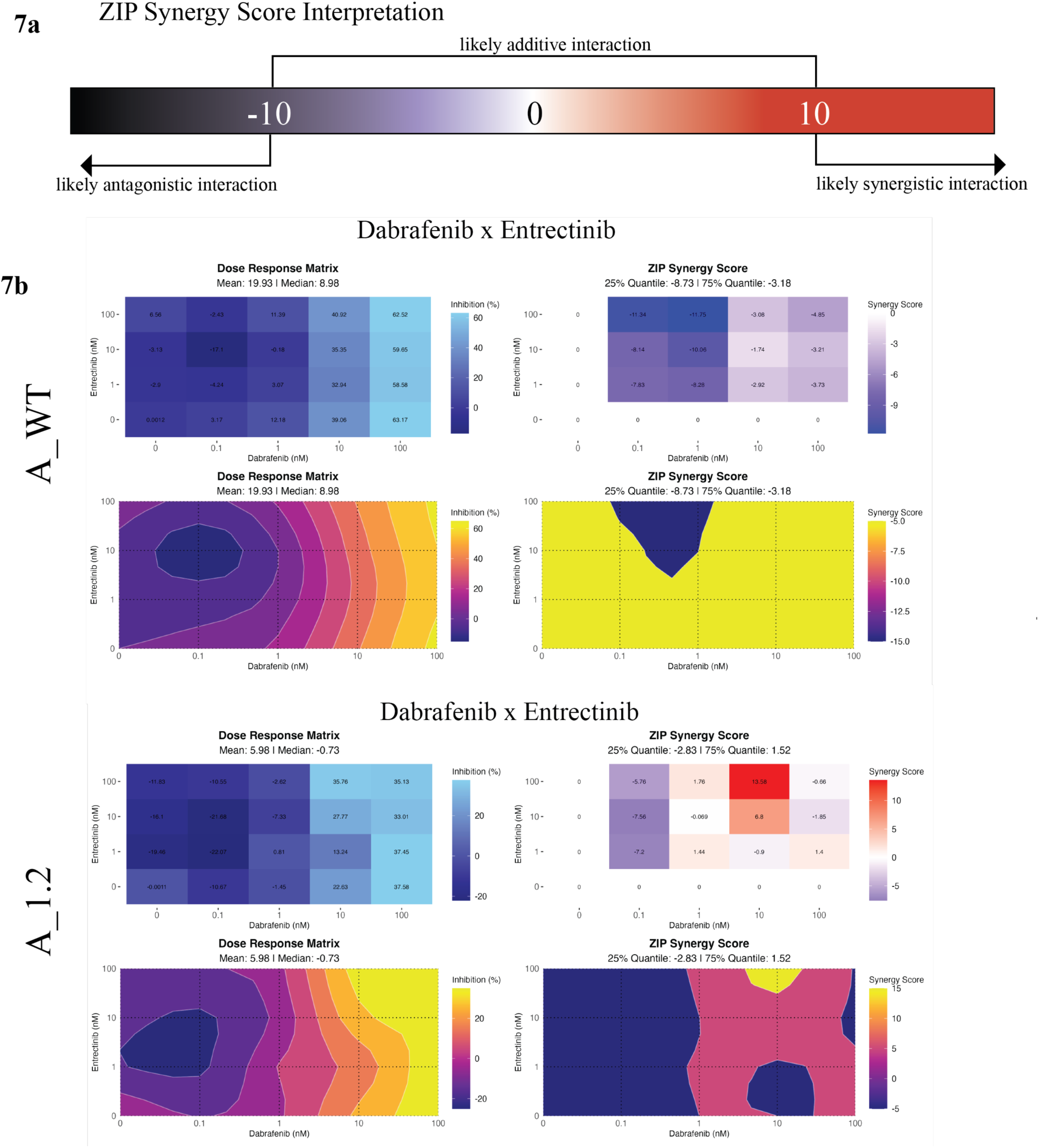
NTRK1 as a potential new treatment option for melanoma 7a. Synergy score range, adapted from SynergyFinder R Package **7b.** Dose-response matrices and ZIP synergy score plots for A_WT and A_1.2 cell lines challenged with combinations of dabrafenib (BRAFi) and entrectonib (NTRKi). Dose-response matrices **(Left):** show the percentage inhibition across a gradient of dabrafenib and entrectonib doses. ZIP synergy score plots **(Right**): demonstrate areas of synergy (positive scores). **(Top):** A_WT exhibits minimal synergy with the combination treatment (ZIP synergy score median =-3.18). **(Bottom):** A_1.2 shows pronounced synergy in some combinations, with a maximum ZIP synergy score of 13.68 (ZIP synergy score median = 1.52).

## Discussion

We successfully adapted our zebrafish-derived enhancer analysis workflow to identify candidate regulatory regions for human *SOX10*. This cross-species approach underscores the utility of zebrafish as a high-throughput model system for functional enhancer analysis despite the common lack of extended stretches of sequence conservation in non-coding regions like enhancers/promoters between zebrafish and humans. By leveraging the zebrafish’s strengths in rapid and scalable assays, our framework can be extended to identify and validate additional human regulatory elements, offering a powerful strategy for uncovering novel enhancers in melanoma biology and other diseases.

Through our enhancer analysis in zebrafish, we identified melanoma-specific regulatory elements that play a crucial role in controlling *sox10* expression, thereby driving melanoma initiation and progression. Deleting a key *sox10* enhancer in zebrafish significantly delayed melanoma onset, highlighting *sox10’s* role in reactivating neural crest transcriptional programs necessary for oncogenic transformation. Translating these findings to human cells, we identified analogous human enhancer regions (Regions 1 and 4) typified by active enhancer chromatin marks (H3K27ac) in melanoma cells at evolutionary conserved (in higher vertebrates) regions with paired *SOX10* binding sites regulating *SOX10* expression. CRISPR-mediated deletion of these now *bona fide* enhancers in melanoma cell lines resulted in lowered *SOX10* expression and slower growth (A375 cells) or more general rewiring of the transcriptome to adapt to *SOX10* loss (WM115 cells) corresponding to the degree of initial SOX10-dependency, confirming their critical role in *SOX10*- driven melanoma.

The use of multiple cell lines with varying baseline phenotypes and degrees of *SOX10* dependency provided a more comprehensive view of transcriptional plasticity under survival pressures. Unlike single cell-line approaches – which may yield a more limited perspective – using a spectrum of genetically engineered enhancer deletion cell lines captured how differential *SOX10* dependency, reminiscent of initial intratumoral heterogeneity, influences cellular responses to selective pressures like enhancer deletion.

Interestingly, some cells with deleted *SOX10* enhancers escaped *SOX10* dependency, shifting toward a more invasive, mesenchymal state – a phenotype commonly linked to drug resistance in melanoma. Bulk RNA sequencing of these phenotype-switched cells revealed a global transition towards an invasive transcriptional profile, closely aligning with drug-resistant subtypes observed in clinical melanoma cases. This transition demonstrated the inherent plasticity of melanoma cells under selective pressure, paralleling both published melanoma sub-phenotype classifications and intratumoral heterogeneity seen in patient samples. These findings emphasize the importance of understanding the poorly characterized drivers of phenotype switching and their contribution to targeted drug resistance.

Further, our study introduces a two-part therapeutic strategy to target melanoma. First, by selectively reducing *SOX10* expression through melanoma-specific enhancer elements; and second, by blocking phenotype switching to prevent drug resistance. Our results suggest that targeting *NTRK1* could provide an additional therapeutic target. *NTRK1* was generally upregulated in most *SOX10* enhancer-deleted cell lines, was well as in drug-resistant melanoma cell lines and human tumors. Given *NTRK1’s* role in activating the MAPK pathway – a central driver of melanoma progression – concurrent inhibition of BRAF/MEK (using dabrafenib/trametinib) and *NTRK1* (using larotrectinib or entrectinib) have additive or even synergistic effects in our cell lines. This multi-pronged-inhibition strategy could block the proliferative MAPK pathway while preventing phenotype switching toward a mesenchymal, drug-resistant state, thus enhancing drug sensitivity in melanoma cells.

*NTRK1*, a neurotrophic tyrosine kinase receptor family member, plays a critical role in activating the MAPK pathway. Although *NTRK* fusions are rare in cutaneous melanoma (<1%), several studies of other tumor types such as infantile fibrosarcoma (*ETV6-NTRK3*), lipofibromatosis-like neural tumor *(LMNA-NTRK1),* low grade spindle cell carcinoma(*RBPMS- NTRK3*), high-grade spindle cell sarcoma (*TMB3-NTRK1*) and fibrohistyocitic proliferation of the skin (*IRF2BP2-NTRK1*), suggest *NTRK1* expression is associated with low or absent *SOX10* expression^70,71^. This supports our hypothesis that reduced *SOX10* dependency drives *NTRK1* upregulation and contributes to phenotype shifts. Furthermore, resistance to *TRK* inhibition has been linked to MAPK pathway reactivation^72^, aligning with our findings that *NTRK1* and *SOX10* act as antagonistic forces in melanoma progression and drug response. Correlations between trajectory position scores and IC50 values for dabrafenib and trametinib further support the link between phenotypic plasticity and drug resistance. Thus, trajectory scoring may serve as a predictive tool for therapeutic response, highlighting the importance of targeting phenotype stability in melanoma treatment.

Additionally, our findings suggest that *NTRK1* inhibition could prevent mesenchymal, drug-resistant phenotype switching, even in lines without overt *NTRK1* overexpression. This highlights its potential as a novel therapeutic target in melanoma and merits future study to fully delineate *NTRK1’s* role in the phenotype regulatory network and drug resistance.

Beyond *NTRK1*, our study identified three additional candidate genes (*AMIGO2, RCOR2,* and *NKX3.1*), each upregulated across most deletion lines and previously reported in lists of upregulated genes in drug-resistant melanoma tumors. *AMIGO2* has been implicated in cell adhesion and tumor progression, both GSEA pathways that were most significantly upregulated in our deletion lines^73–75^. Neural development regulator *RCOR2* has also been associated with transcriptional reprogramming in glioblastoma, another potentially neural crest derived solid tumor^76^. Finally, loss of *NKX3.1,* a critical factor for prostate cancer cell differentiation, with emerging evidence of roles in regulating transcriptional plasticity, was also studied^77^. Of these prioritized genes, *RCOR2* and *NTRK1* siRNA knockdown yielded the most striking results in terms of restoring drug sensitivity in human melanoma cell lines with SOX10 enhancer deletions studied here.

Future research will focus on further elucidating *NTRK1’s* role within the phenotype regulatory network and exploring how it intersects with *SOX10*-dependent pathways. Clinical trials combining BRAF/MEK inhibitors with *NTRK* inhibitors could validate the effectiveness of targeting these pathways simultaneously. By addressing both upstream and downstream components of the *SOX10* regulatory axis, our findings pave the way for novel therapeutic strategies against melanoma plasticity and drug resistance.

## Acknowledgments

We thank Rebecca Cunningham for helpful discussions and comments on the manuscript. The content is solely the responsibility of the authors and does not necessarily represent the official views of the NIH. C.K.K. was funded by the Cancer Research Foundation Young Investigator Award and NIH R01CA240633. SND was funded by the National Science Foundation Graduate Research Fellowship Program and the Cellular and Molecular Biology T32 Training Grant at Washington University in St. Louis. Research reported in this publication was supported in part by the National Cancer Institute of the National Institutes of Health (NIH) under award number R01CA240633. We thank the Genome Engineering & Stem Cell Center at Washington University for assistance with cell line generation. We thank the Genome Technology Access Center in the Department of Genetics at Washington University School of Medicine for performing deep sequencing. The Center is partially supported by NCI Cancer Center Support Grant #P30 CA91842 to the Siteman Cancer Center and by ICTS/CTSA Grant# UL1 TR000448 from the National Center for Research Resources (NCRR), a component of the National Institutes of Health (NIH), and NIH Roadmap for Medical Research.

## Methods and Materials

### Chromatin Immunoprecipitation Sequencing (ChIP-seq) Analysis

To investigate enhancer regions upstream of the *SOX10* locus, we analyzed ChIP-seq data from Kaufman^1^ using the UCSC Genome Browser. H3K27ac tracks were visualized for multiple human cell lines, including CJM, COLO679, SKMEL2, SKMEL30, UAC257, A375 and a NCC cell line. A ∼60 kb stretch of H3K27ac peaks consistently observed across several melanoma cell lines was identified, starting approximately 30 kb upstream of the *SOX10* transcription start site.

### Conservation Analysis

Regions with evolutionary conservation were identified using the “Vertebrate Conservation” tracks in the UCSC Genome Browser, followed by confirmation with the ECR Browser. Conservation scores were retrieved from the hg19 100-way PhastCons dataset. A genomic region encompassing the *SOX10* locus (chr22:38380408-38449849, hg19) was defined as a GRanges object. Conservation scores within this region were imported from the PhastCons bigwig file. The scores were analyzed and visualized to identify patterns of high conservation. Genomic coordinates were divided into 1000bp tiles, and average PhastCons scores were calculated for each tile. High-conservation tiles, defined as those with average scores >0.75, were further analyzed.

### Motif Scanning in High-Conservation Regions

Sequences for high conservation 1000 bp tiles were extracted from the hg19 genome using the BSgenome.Hsapiens.UCSC.hg19 package. Motif scanning was performed using the motifmatchr package and SOX10 transcription factor binding motifs from JASPAR 2022 (motif IDs MA0442.1 and MA0442.2). Matches were identified for both motifs, and sequences containing these matches were extracted. We then selected/focused our attention further on Region 1 and 4 for further analysis based on published SOX10 ChIP-seq^46^ showing *bona fide* SOX10 binding and suggestive evidence of enhancer function in reporter screen context^45^.

A375 and WM115 cell lines were chosen after consulting the DepMap CERES Gene Effect dataset, accessed on 5/20/21.

### Zebrafish lines and rearing conditions

Zebrafish were bred and maintained following Washington University IACUC animal care protocols. Adult fish were bred as either pairs or groups, and resulting embryos were reared in egg water (5 mM NaCl, 0.17 mM KCl, 0.33 mM CaCl2_22, 0.33 mM MgSO4_44) at 28.5°C. The study utilized the following zebrafish strains and transgenic lines:

*Tg(BRAF^V600E^);p53^−/−^;peak5^stl5^*^38^

*Tg(BRAF^V600E^);p53^−/−^*.

### Zebrafish reporters of human enhancer function

For cloning of regions of interest, genomic DNA was extracted from A375 and WM115 human melanoma cells using the GenElute Mammalian Genomic DNA Miniprep Kit using manufacturer instructions. The regions of interest were initially amplified from this genomic DNA using Phusion polymerase (Table 1, PCR primer sequences), then gel extracted with a QIAquick Gel Extraction Kit.

The BFMP-FK_EGFP plasmid was used as previously described^3,40^, and mutated to add a SalI restriction enzyme site via Q5 Mutagenesis. After linearization, the putative enhancer regions “Region 1” and “Region 4” were inserted into the backbone vector using NEB HiFi Assembly (see **Supplementary Table 7** for HiFi primers), and then transformed into TOP10 cells. Whole-plasmid sequencing confirmed successful integration.

For analysis of enhancer function, GFP reporter plasmid was co-injected with a similar Tol2 plasmid containing the zebrafish *Sox10* promoter driving mCherry (concentration of DNA and Tol2) per standard Tol2-based transgenesis approaches^78^. Resulting F0 embryos were screened for fluorescence on days 1-5 dpf using a Nikon SMZ18 fluorescent dissecting microscope under the long pass and short pass filters to assess GFP and mCherry activity and localization.

### Cell Culture

A375 human melanoma cells (acquired from ATCC) were maintained in DMEM with 10% FBS and 1% P/S. WM115 cells were purchased from Fisher Scientific (NC1926427) and were maintained in Tu2% medium, prepared as follows: 1 L of MCDB medium was prepared by dissolving 1 bottle of MCDB (cat # M74031L) in 1 L of ddH20 with 1.2 g of sodium bicarbonate. To this, 250 mL of L-15 medium (cat # 11415114), 25 mL of FBS (cat # A3160601), 1.25 mL of insulin (cat # I0516-5ML), 1.5 mL of calcium chloride (cat # BP974210X5), and 12.78 mL of P/S were added, as previously described on the Herlyn Lab website. The solution was mixed thoroughly and used to culture WM115 cells. All cells were maintained at in a 37 incubator at 5% CO2.

### Generation of targeted genomic deletion cell lines

To generate cell lines with targeted deletions upstream of human SOX10 (Region 1 and Region 4), we collaborated with the Genome Engineering and Stem Cell Center (GESC) core at Washington University (https://geneediting.wustl.edu/).

The GESC core designed gRNAs using a CRISPR algorithm to minimize off target effects. These gRNAs were synthesized as sgRNAs by IDT (**Supplementary Table 8**). WM115 and A375 cells were trypsinized, counted (100,000-200,000 cells per reaction), and electroporated with ribonucleoprotein complexes of Cas9 and sgRNAs targeting Region 1 or Region 4. Transfected cells were seeded in pools, allowed to recover for 72 hours, and subsequently genotyped by PCR to confirm deletions within the targeted regions. PCR screening included both the cutting sites and a 300 bp window across the expected deletion region.

If deletions were detected, single cells were sorted into 96-well plates using a Sony SH800 fluorescent cell sorter. Clonal populations were expanded and re-genotyped. This process yielded nine deletion lines and four WT lines across both cell types. WT cells from A375 and WM115 were also subjected to single-cell sorting and clonal expansion to mimic the bottleneck experienced by the deletion lines.

In A375 cells, two stable deletion lines were generated for Region 1 (A_1.1, A_1.2) and two for Region 4 (A_4.1, A_4.2). For WM115 cells, three deletion lines were established for Region 1 (W_1.1, W_1.2, W_1.3) and two for Region 4 (W_4.1, W_4.2).

### Western blot analysis

Proteins were extracted from each deletion line and their corresponding WT lines using a PMSF-containing cell lysis buffer. The lysate was passed through a 21G needle to ensure homogenization, and the protein extracts were stored at-20°C. Western blotting was performed using Criterion XT precast gels (Ref #1610374), following previously established protocols.

The following antibodies and reagents were used:

- Primary antibody: SOX10 (ab227680, rabbit monoclonal antibody, 1:400 dilution)
- Loading control: GAPDH (14C10 rabbit monoclonal antibody, #2118)
- Secondary antibody: LICOR IRDye 680RD goat anti-rabbit IgG
- Protein ladder: Precision Plus Protein Kaleidoscope Prestained Protein Standards (Cat #1610375)

### Cell proliferation assays

WM115 cells were seeded at 1000 cells/well, and A375 cells were seeded at 500 cells/well in opaque walled 96 well plates (Corning Ref#3903). Cell proliferation was quantified using the Cell Titer Glo 2.0 kit, and measured using a luminometer on Day 1, 3, 5, and 7. Dunnett’s multiple comparisons test was used to compare the WT to the experimental conditions.

### RNA-seq analysis

RNA was extracted from each melanoma line, included engineered cell lines, and corresponding WT lines using the Qiagen RNeasy Mini Kit (Ref # 74104) and Qiagen RNeasy Plus Mini Kit (Ref # 74034). At least three separate samples of RNA were taken from different time points and different passage numbers for each deletion line. RNA quality was assessed using the nanodrop, and only samples with concentrations >200 ng/uL and the appropriate 260/280 and 260/230 ratios were used in future experiments. RNA was stored at-80. cDNA was generated using the SuperScript III RT kit (Ref # 12574026).

Multiple unique samples of RNA from each deletion line and corresponding WT lines were submitted to the Genetics GTAC. This core assessed the quality of the submitted RNA and the three best unique samples (RIN = 10) from each line were used moving forward. Genetics GTAC constructed a sequencing library and performed RNA-sequencing on the Illumina NovaSeq6000 S4 XP flow cells with 2×150 paired-end reads.

### Phenotype Score Calculation

Phenotype scores were calculated as previously described by Tsoi^19^. Briefly, sample counts were imported into R and filtered to include only genes belonging to the Tsoi subtype groups. For each cell line, the mean expression value was calculated within each sub-phenotype category. Z-scores were then computed for these subtype scores. To derive a weighted identity score, the Z-scores were normalized on a scale of 0-1, and each category was multiplied by its respective weight (the category number divided by the sum of all unweighted identity scores).

### Pathway enrichment analysis

Pathway enrichment analysis was run in R using gene sets from: https://www.gsea-msigdb.org/gsea/msigdb/index.jsp

Gene sets used for GSEA analysis, downloaded from gsea-msigdb.org:

- C2: Curated Gene sets (7233 gene sets)

`o` Canonical pathways (3795 gene sets)
- C3: Regulatory Gene Sets (3713 gene sets)
- C4: Computational Gene Sets (1007 gene sets)
- C5: Ontology Gene Sets (16008 gene sets)
- C6: Oncogenic Gene Sets (189 gene sets)
- C8: Cell Type Signature Gene Sets (830 gene sets)
- H: Hallmark Gene Sets (50 gene sets)

GSEA was performed using the sorted gene lists for each deletion line with the fgsea package in R. For each deletion line, the top 10 most upregulated and top 10 most downregulated gene sets from each collection were extracted and saved into a new dataframe. Dataframes from all deletion lines were then joined based on shared differentially expressed gene sets. Gene sets were ranked in descending order by the number of deletion lines in which they appeared as a top 10 hit.

### Drug Sensitivity Assays

Melanoma cells were plated in clear bottomed 96 well plates at a seeding density of 1.5E3 cells per well. Drugs (Dabrafenib: Fisher Scientific NC0621920, and Trametinib: Fisher Scientific NC2307384) were serially diluted in sterile DMSO and stored per the manufacturers instructions. 24 hours after seeding, media was replaced with media containing serially diluted drugs at noted concentrations. Vehicle wells were given media with the same volume of plain DMSO. Cells were incubated at 37 for 72 hours, after which cell proliferation was quantified using the Cell Titer Glo 2.0 kit, and measured using a luminometer.

### siRNA Knockdown of Target Genes

The following siRNA products were used:

Negativeive CTL (catalog number: AM4611), 5 nmol

NTRK1 (AM16708), 5 nmol

NKX3.1 (AM16708), 5 nmol

RCOR2 (AM16708), 5 nmol

AMIGO2 (AM16708), 5 nmol

Cells were plated at a seeding density of 1.5E3 cells per well in antibiotic free media. Cells were allowed to grow to 60-80% confluence, after which siRNA and lipofectamine were added. After a 24 hour incubation, the media was replaced with media containing but siRNA and a standard dilution curve concentration of either Dabrafenib or Trametinib. Vehicle wells were treated with siRNA but received DMSO in place of drug. Cells were then incubated at 37 for an additional 48 hours, after which cell proliferation and survival was quantified using Cell Titer Glo 2.0 kit, measured using a luminometer, as described above.

### Data availability

RNA-seq data available through GEO, series number: GSE283223.

### Code availability

All scripts and code used to generate the results and figures in this paper are available at the GitHub repository: https://github.com/nofedege/Melanoma_Drug_Resistance_Data.

## Notes

**Conflict of Interest Statement:** Authors declare no competing interests

### Competing Interest Statement

The authors have declared no competing interest.

