## Supplementary material for "Specific *SOX10* enhancer elements modulate phenotype plasticity and drug resistance in melanoma": Supp Figures and Tables

**Supplemental Figure 1. Identification and Functional Testing of Melanoma-Associated Enhancers in Zebrafish**

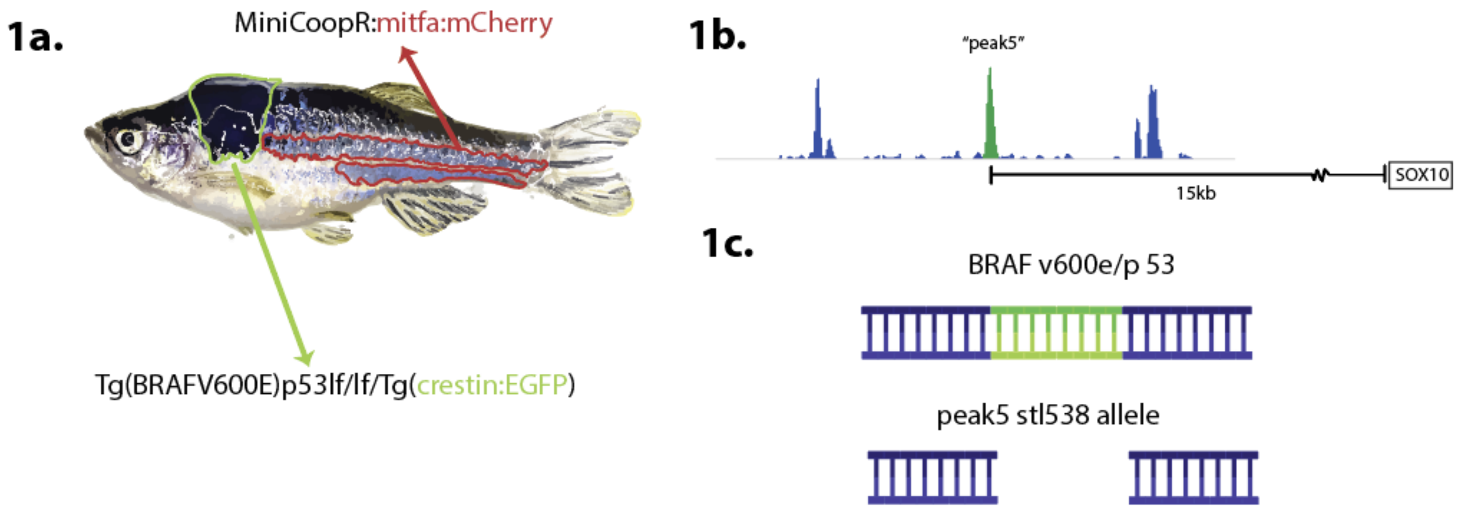

**Supplemental Figure 1:**
**1a.** Cells from zebrafish melanoma tumors, marked by Tg(BRAFV600E)/ p53lf/lf/ Tg(crestin:EGFP) were sorted to separate them from other cell populations. Melanocytes were marked specifically by MiniCoopR:mitfa:mCherry. **1b.** ATAC-seq was performed on isolated tumor cells to identify regions of differentially accessible chromatin (i.e. more open in melanoma than melanocytes), suggesting putative enhancer elements specific to melanoma (Kramer 2021). One melanoma-specific peak, termed "peak5," was identified as a candidate enhancer **1c.** stl538 deletion line, generated by Cunninham et al. 2021, involves a CRISPR Cas9-mediated deletion of a conserved region within the ATAC-seq identified "peak5" enhancer element of the *sox10* locus (Cunningham 2021) .

**Supplemental Figure 2. Comparative Workflow for Identifying Candidate Regulatory Regions in Human Melanoma from Zebrafish**

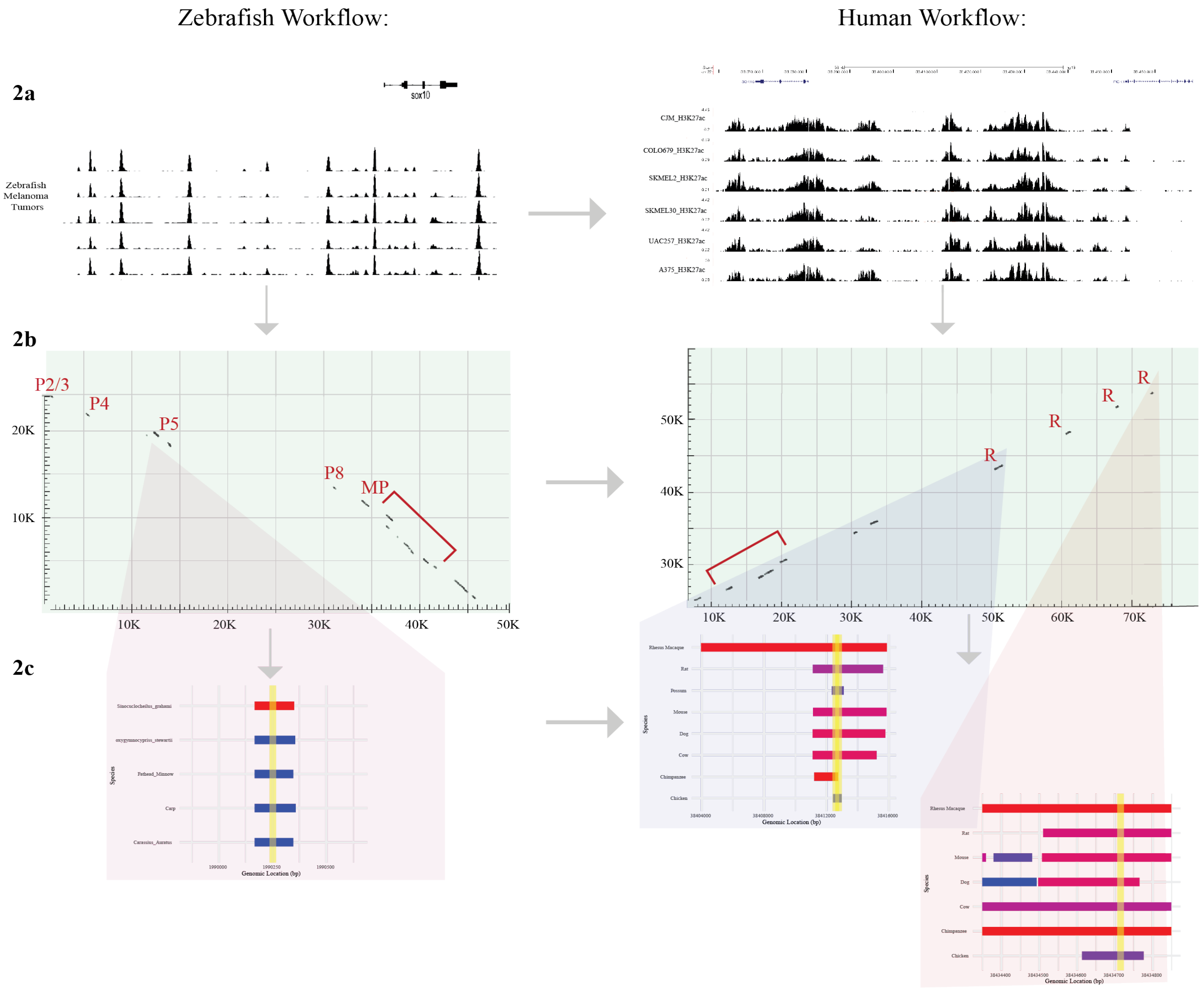

**Supplemental Figure 2:**
**2a.** Initial identification of candidate regulatory regions based on differential chromatin accessibility in melanoma cells versus wild-type melanocytes. **(Left):** Zebrafish ATAC-seq peaks highlighting regions of accessible chromatin in melanoma cells (adapted from Cunningham et al.). **(Right):** H3K27ac ChIP-seq profiles in human melanoma cell lines, indicating active chromatin regions (adapted from Kaufman et al.) **2b.** Further refinement of candidate regions by evolutionary conservation. Sequence conservation marks in dot plots suggest functional significance. **Left:** Conservation between zebrafish and Cyprinidae family member carp, with conserved sequences aligning with accessible chromatin peaks. **Right:** Conservation between human and rat, with conserved regions aligning with four major H3K27ac peaks. **2c.** Examination of conserved regions for the presence of SOXE dimer binding sites. **Left:** Alignment of zebrafish "peak5" with other Cyprinidae genomes reveals a conserved SOXE dimer site. **Right:** Only two conserved regions in human alignments show potential for a SOXE dimer binding site (highlighted in yellow).

**Supplemental Figure 3**: **Western blot analysis of SOX10 expression in A375 and WM115 melanoma cell lines.**

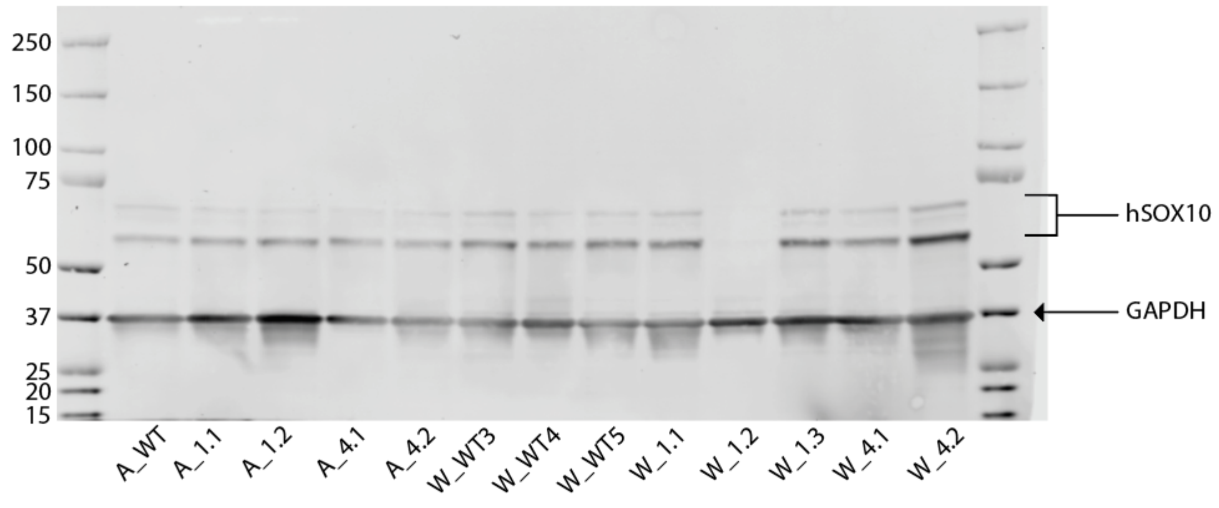

**Supplemental Figure 3**: Western blot analysis of SOX10 expression in A375 and WM115 melanoma cell lines. Protein lysates from A375 wild-type (WT) and mutant lines (A_1.1, A_1.2, A_4.1, A_4.2) and WM115 wild-type (WT3, WT4, WT5) and mutant lines (W_1.1, W_1.2, W_1.3, W_4.1, W_4.2) were probed with antibodies against SOX10 and GAPDH (loading control). The blot shows a decrease in SOX10 expression in specific mutant lines compared to their respective wild-type controls, highlighting potential functional consequences of genetic or regulatory perturbations in these models.

**Supplemental Figure 4: Integrated analysis of gene expression and pathway enrichment across A375 and WM115 enhancer-deleted melanoma cell lines.**

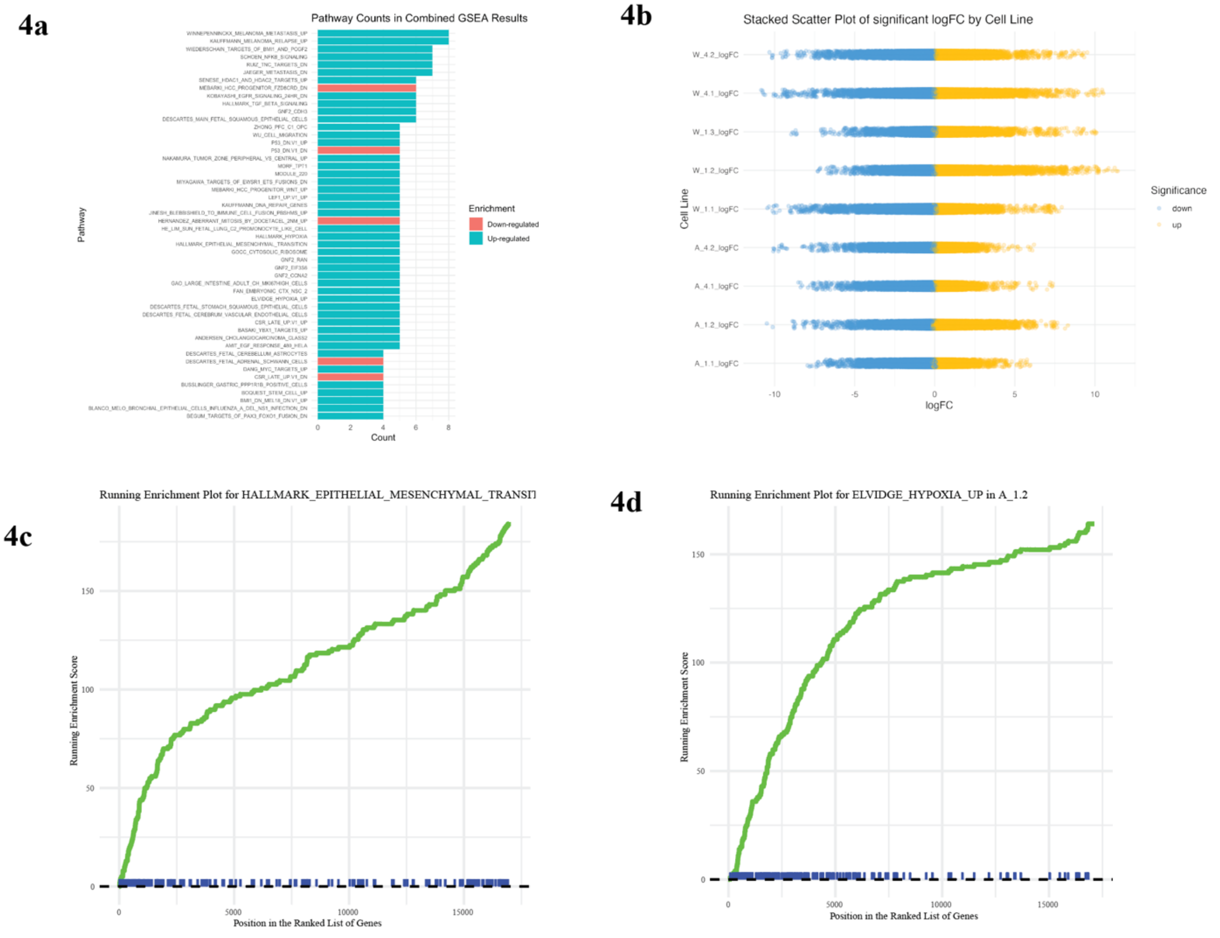

**Supplemental Figure 4:**

**4a:** Bar plot displaying pathway counts from combined GSEA results. Pathways are categorized as upregulated (teal) or downregulated (red) based on enrichment scores. Key pathways include epithelial-mesenchymal transition (EMT) and hypoxia, suggesting significant shifts in cell phenotypes due to enhancer deletions. **4b:** Stacked scatter plot of significant log fold-change (logFC) values by cell line. Each point represents a gene with significant differential expression (colored by enrichment category). Mutant lines exhibit both upregulation and downregulation patterns compared to wild-type, reflecting transcriptional plasticity. **4c:** Running enrichment plots for representative pathways. The left panel shows enrichment for the hallmark epithelial-mesenchymal transition (EMT) pathway, while the right panel (**4d)** highlights the Elvidge hypoxia response pathway. Both demonstrate substantial enrichment in mutant line A_1.2, emphasizing the role of deleted enhancers in modulating stress response and phenotypic plasticity.

**Supplemental Figure 5: MAPKi Resistance Up-Regulated Genes**

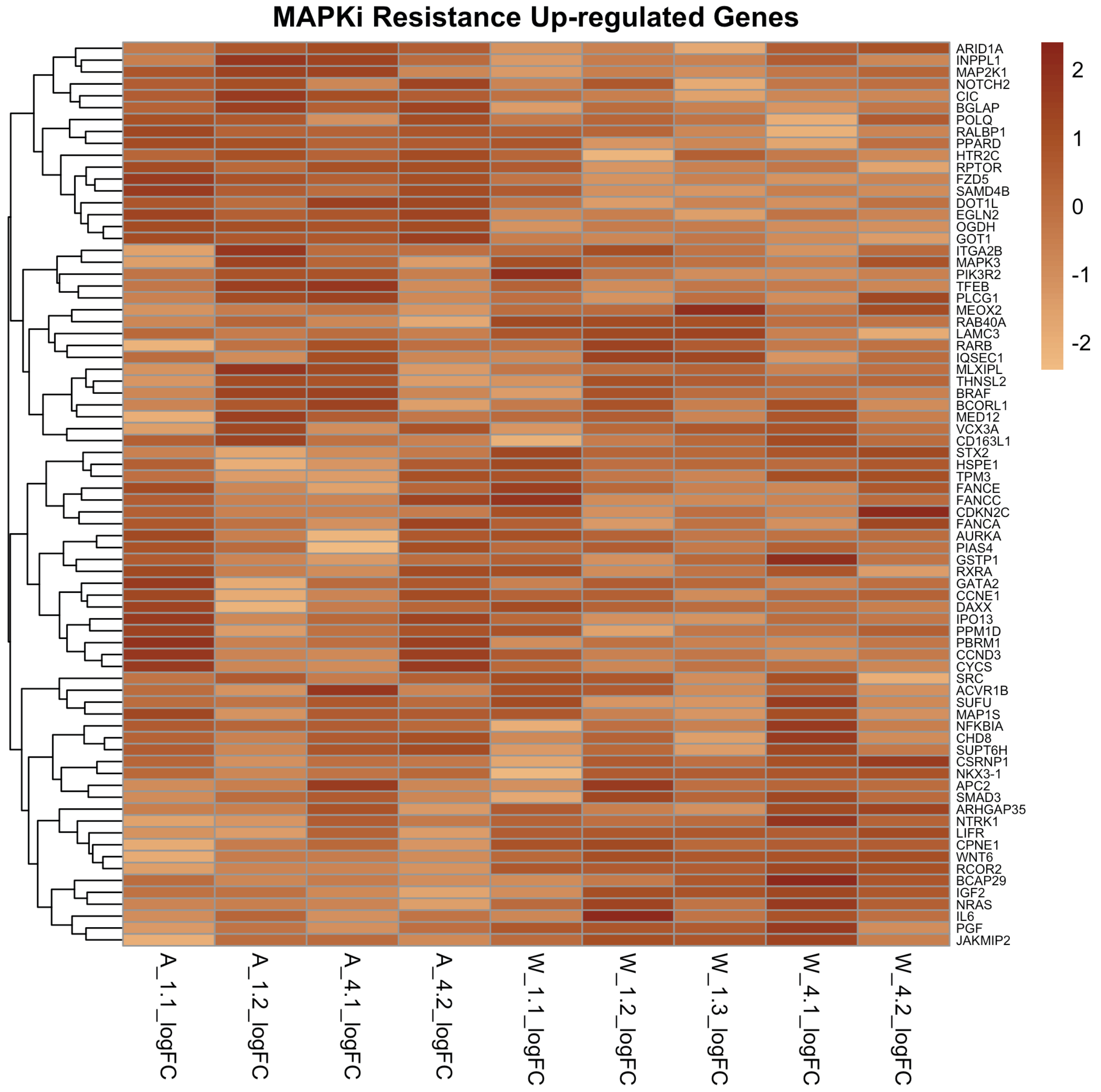

**Supplemental Figure 5:**

Log2 fold-change (logFC) values for all deletion lines in comparison to the top genes in the MAPK inhibitor (MAPKi) resistance upregulated gene set reported by Hugo et al., 2015. Genes are hierarchically clustered to illustrate patterns of expression.

**Supplementary Table 1. Comparative genomic alignments of conserved regions upstream of *SOX10* (Regions 1 and 4) across species**

| REGION | SPECIES | COMPARATIVE GENOME | ALIGNMENT BLOCK OTHER SPECIES | ALIGNMENT BLOCK HUMAN | IDENTITY MATCH PERCENT | LENGTH BPS |
|---|---|---|---|---|---|---|
| **REGION 1** | Chimpanzee | panTro3 | chr22:36666061-36667598 | chr22:38411208-38412746 | 100.0% | 260 |
| **REGION 1** | Possum | monDom5 | chrUn:100678219-100679011 | chr22:38412487-38412850 | 77.7% | 364 |
| **REGION 1** | Possum | monDom5 | chrUn:100678219-100679011 | chr22:38412341-38413111 | 80.2% | 86 |
| **REGION 1** | Chicken | galGal3 | chr1:52992159-52992805 | chr22:38412422-38412982 | 77.8% | 424 |
| **REGION 1** | Rat | rn4 | chr7:117174967-117178989 | chr22:38411121-38415635 | 86.0% | 501 |
| **REGION 1** | Mouse | mm10 | chr15:79191984-79196126 | chr22:38411134-38415855 | 90.8% | 501 |
| **REGION 1** | Cow | bosTau6 | chr5:110306361-110310536 | chr22:38411114-38415219 | 92.8% | 501 |
| **REGION 1** | Dog | canFam2 | chr10:29722656-29727387 | chr22:38411114-38415775 | 93.0% | 501 |
| **REGION 1** | Rhesus Macaque | rheMac2 | chr10:81941996-81953607 | chr22:38403951-38415877 | 98.2% | 501 |
| **REGION 1** | Chimpanzee | panTro3 | chr22:36666061-36667598 | chr22:38411208-38412746 | 100.0% | 260 |
| **REGION 1** | Possum | monDom5 | chrUn:100678219-100679011 | chr22:38412487-38412850 | 77.7% | 364 |
| **REGION 1** | Possum | monDom5 | chrUn:100678219-100679011 | chr22:38412341-38413111 | 80.2% | 86 |
| **REGION 1** | Chicken | galGal3 | chr1:52992159-52992805 | chr22:38412422-38412982 | 77.8% | 424 |

| REGION | Species | Comparative Genome | Alignment Block other species | Alignment Block Human | Identity Match Percent | Length bps |
|---|---|---|---|---|---|---|
| **REGION 4** | Chicken | galGal3 | chr1:52980839-52981002 | chr22:38434613-38434776 | 0.73 | 164 |
| **REGION 4** | Dog | canFam2 | chr10:29706729-29706873 | chr22:38434349-38434493 | 0.67 | 145 |
| **REGION 4** | Dog | canFam2 | chr10:29706452-29706728 | chr22:38434497-38434765 | 0.87 | 269 |
| **REGION 4** | Rat | rn4 | chr7:117196679-117197039 | chr22:38434510-38434849 | 0.85 | 340 |
| **REGION 4** | Mouse | mm10 | chr15:79212787-79213157 | chr22:38434507-38434849 | 0.87 | 343 |
| **REGION 4** | Mouse | mm10 | chr15:79212656-79212763 | chr22:38434379-38434481 | 0.70 | 103 |
| **REGION 4** | Mouse | mm10 | chr15:79212633-79212646 | chr22:38434349-38434359 | 0.82 | 11 |
| **REGION 4** | Cow | bosTau6 | chr5:110328308-110328831 | chr22:38434349-38434849 | 0.80 | 501 |
| **REGION 4** | Rhesus Macaque | rheMac2 | chr10:81971709-81972213 | chr22:38434349-38434849 | 0.96 | 501 |
| **REGION 4** | Chimpanzee | panTro3 | chr22:36690339-36690838 | chr22:38434349-38434849 | 0.98 | 501 |
| **REGION 4** | Chicken | galGal3 | chr1:52980839-52981002 | chr22:38434613-38434776 | 0.73 | 164 |

**Supplementary Table 1. Comparative genomic alignments of conserved regions upstream of *SOX10* (Regions 1 and 4) across species**
Alignment details of Regions 1 and 4 across various species, including their respective comparative genome assemblies, alignment blocks in other species, and their corresponding alignment blocks in the human genome (hg19). Identity match percentage represents the level of nucleotide conservation between the aligned regions, and length in base pairs (bps) specifies the size of the aligned block.

Species listed include primates (e.g., chimpanzee, rhesus macaque), mammals (e.g., possum, rat, mouse, cow, dog), and birds (e.g., chicken). Data highlights conserved sequences between human and other vertebrates, underscoring the evolutionary conservation of these regions.

**Supplementary Table 2: Correlation of growth rate and SOX10 expression levels across WM115 lines**

| Cell Line | Growth rate (*k*) | $R^2$ of *k* | Average $2^{-\Delta Ct}$ SOX10 expression | SD of SOX10 expression |
|---|---|---|---|---|
| **W_WT** | 0.1727 | 0.9831 | 0.09746248 | 0.01490698 |
| **W_1.1** | 0.2076 | 0.9925 | 0.04627999 | 0.00667862 |
| **W_1.2** | 0.3364 | 0.9605 | 0.0036405 | 0.00438119 |
| **W_1.3** | 0.2239 | 0.9819 | 0.06470216 | 0.04387434 |
| **W_4.1** | 0.2413 | 0.9949 | 0.05368586 | 0.00667862 |
| **W_4.2** | 0.2413 | 0.9917 | 0.0284234 | 0.0084183 |

| | |
|---|---|
| **Pearson r** | -0.8909 |
| **R squared** | 0.7937 |
| **P value (two-tailed)** | 0.0172 |
| **Significance (alpha = 0.05)** | Yes: * |

**Supplementary Table 2:** Growth rates (*k*), goodness of fit ($R^2$), and SOX10 expression levels for all WM115 cell lines. Growth ratios were calculated using normalized proliferation data from CellTiter-Glo assays. Proliferation was normalized to the Day 1 mean for each cell line, and exponential growth rates (*k*) were determined by comparing values from Day 1 to Day 5. SOX10 expression was measured using qPCR and is presented as the average $2^{-\Delta Ct}$ values with standard deviations (SD). The correlation between growth rate (*k*) and SOX10 expression was calculated using Pearson's correlation ($r = -0.8909$, $p = 0.0172$), showing a statistically significant negative relationship (*, $p<0.05$).

**Supplementary Table 3. Phenotype trajectory scores and statistical comparisons for deletion and WT lines.**

| | A_WT | A_1.1 | A_1.2 | A_4.1 | A_4.2 | W_WT | W_1.1 | W_1.2 | W_1.3 | W_4.1 | W_4.2 |
|---|---|---|---|---|---|---|---|---|---|---|---|
| **MEAN** | 4.4310 | 4.74980 | 4.97530 | 4.38540 | 4.66829 | 3.56896 | 3.60807 | 4.43976 | 3.81301 | 4.38676 | 4.34717 |
| **SD** | 0.1463 | 0.28989 | 0.13863 | 0.06895 | 0.14841 | 0.25068 | 0.14635 | 0.49245 | 0.46277 | 0.07551 | 0.06586 |
| **N** | 4 | 3 | 4 | 4 | 3 | 4 | 5 | 3 | 3 | 3 | 3 |

| DUNNETT'S MULTIPLE COMPARISONS TEST | MEAN DIFF. | 95.00% CI OF DIFF. | BELOW THRESHOLD? | SUMMARY | ADJUSTED P VALUE |
|---|---|---|---|---|---|
| **W_1.3 VS. W_WT5** | 0.2441 | -0.3352 to 0.8233 | No | ns | 0.7379 |
| **W_1.2 VS. W_WT5** | 0.8708 | 0.2915 to 1.450 | Yes | ** | 0.0021 |
| **W_1.1 VS. W_WT5** | 0.03911 | -0.4697 to 0.5479 | No | ns | >0.9999 |
| **W_4.2 VS. W_WT5** | 0.7782 | 0.1990 to 1.357 | Yes | ** | 0.0059 |
| **W_4.1 VS. W_WT5** | 0.8178 | 0.2385 to 1.397 | Yes | ** | 0.0038 |
| **W_WT4 VS. W_WT5** | 0.4042 | -0.1750 to 0.9835 | No | ns | 0.2579 |
| **W_WT3 VS. W_WT5** | 0.2548 | -0.3244 to 0.8341 | No | ns | 0.7022 |
| **W_1.3 VS. W_WT5** | 0.2441 | -0.3352 to 0.8233 | No | ns | 0.7379 |
| **W_1.2 VS. W_WT5** | 0.8708 | 0.2915 to 1.450 | Yes | ** | 0.0021 |
| **W_1.1 VS. W_WT5** | 0.03911 | -0.4697 to 0.5479 | No | ns | >0.9999 |
| **A_4.2 VS. A_WT** | 0.2372 | -0.1113 to 0.5858 | No | ns | 0.2313 |
| **A_4.1 VS. A_WT** | -0.04568 | -0.3684 to 0.2770 | No | ns | 0.9846 |
| **A_1.2 VS. A_WT** | 0.5442 | 0.2215 to 0.8669 | Yes | ** | 0.0015 |
| **A_1.1 VS. A_WT** | 0.3187 | -0.0298 to 0.6673 | No | ns | 0.0769 |
| **A_4.2 VS. A_WT** | 0.2372 | -0.1113 to 0.5858 | No | ns | 0.2313 |
| **A_4.1 VS. A_WT** | -0.04568 | -0.3684 to 0.2770 | No | ns | 0.9846 |

| ANOVA summary: A375 Lines | |
|---|---|
| **F** | 6.315 |
| **P value** | 0.0006 |
| **P value summary** | *** |
| **Significant diff. among means (P < 0.05)?** | Yes |
| **R squared** | 0.6994 |

| ANOVA summary: WM115 Lines | |
|---|---|
| **F** | 8.6 |
| **P value** | 0.0013 |
| **P value summary** | ** |
| **Significant diff. among means (P < 0.05)?** | Yes |
| **R squared** | 0.7257 |

**Supplementary Table 3. Phenotype trajectory scores and statistical comparisons for deletion and WT lines:** Mean phenotype trajectory scores, standard deviations (SD), and sample sizes (N) for each deletion and WT line, highlighting shifts in phenotype identity across cell lines. Scores are derived from weighted trajectory calculations based on Tsoi sub-phenotype gene lists, ranging from 1 (melanocytic) to 7 (undifferentiated). Dunnett's multiple comparisons test results include mean differences, 95% confidence intervals (CI), significance summaries, and adjusted p-values for pairwise comparisons between deletion and WT lines. ANOVA summaries for A375 and WM115 lines indicate significant differences among means, with F-statistics, p-values, and R-squared values provided. These data support the trend toward a more undifferentiated phenotype in SOX10 enhancer-deleted lines.

**Supplementary Table 4. IC50 and dose-response data for WM115 and A375 cell lines treated with dabrafenib.**

| WM115 Dabrafenib | | | | | | |
|---|---|---|---|---|---|---|
| | W_WT | W_1.1 | W_1.2 | W_1.3 | W_4.1 | W_4.2 |
| **log(inhibitor) vs. normalized response** | | | | | | |
| **Best-fit values** | | | | | | |
| **LogIC50** | -0.2523 | -0.4952 | 1.893 | 0.04641 | 1.188 | 0.6152 |
| **IC50** | 0.5594 | 0.3197 | 78.24 | 1.113 | 15.4 | 4.123 |
| **95% CI (profile likelihood)** | | | | | | |
| **LogIC50** | -0.5726 to 0.05597 | -0.9624 to -0.009382 | 1.215 to 2.548 | -0.3628 to 0.5144 | 0.6081 to 1.997 | 0.04424 to 1.173 |
| **IC50** | 0.2676 to 1.138 | 0.1091 to 0.9786 | 16.42 to 353.5 | 0.4337 to 3.269 | 4.056 to 99.20 | 1.107 to 14.91 |
| **Goodness of Fit** | | | | | | |
| **Degrees of Freedom** | 17 | 17 | 17 | 17 | 17 | 17 |
| **R squared** | 0.8337 | 0.658 | 0.4217 | 0.6477 | 0.4829 | 0.6235 |
| **Sum of Squares** | 3144 | 5199 | 6962 | 5601 | 7432 | 6300 |
| **Sy.x** | 13.6 | 17.49 | 20.24 | 18.15 | 20.91 | 19.25 |
| **Number of points** | | | | | | |
| **# of X values** | 18 | 18 | 18 | 18 | 18 | 18 |
| **# Y values analyzed** | 18 | 18 | 18 | 18 | 18 | 18 |

| A375 Dabrafenib | | | | | |
|---|---|---|---|---|---|
| | A_WT | A_1.1 | A_1.2 | A_4.1 | A_4.2 |
| **log(inhibitor) vs. normalized** | | | | | |

| | | | | | |
|---|---|---|---|---|---|
| **response** | | | | | |
| **Best-fit values** | | | | | |
| **LogIC50** | 0.8214 | 0.7924 | 1.034 | 1.224 | 1.283 |
| **IC50** | 6.628 | 6.2 | 10.82 | 16.75 | 19.18 |
| **95% CI (profile likelihood)** | | | | | |
| **LogIC50** | 0.5623 to 1.081 | 0.5394 to 1.038 | 0.6353 to 1.499 | 0.7936 to 1.789 | 0.9429 to 1.682 |
| **IC50** | 3.650 to 12.04 | 3.463 to 10.92 | 4.318 to 31.54 | 6.217 to 61.56 | 8.768 to 48.03 |
| **Goodness of Fit** | | | | | |
| **Degrees of Freedom** | 17 | 17 | 17 | 17 | 17 |
| **R squared** | 0.8827 | 0.903 | 0.7145 | 0.7526 | 0.8353 |
| **Sum of Squares** | 2533 | 2069 | 4463 | 6231 | 4825 |
| **Sy.x** | 12.21 | 11.03 | 16.2 | 19.15 | 16.85 |
| **Number of points** | | | | | |
| **# of X values** | 18 | 18 | 18 | 18 | 18 |
| **# Y values analyzed** | 18 | 18 | 18 | 18 | 18 |

**Supplementary Table 4. IC50 and dose-response data for WM115 and A375 cell lines treated with dabrafenib.** This table provides the results of dose-response experiments for WM115 and A375 cell lines, including wild-type (WT) and SOX10 enhancer deletion lines, treated with the BRAF inhibitor dabrafenib. Best-fit values for log(IC50) and IC50 are reported alongside 95% confidence intervals (CI). Goodness-of-fit metrics include R-squared values, degrees of freedom, and sums of squares. The total number of data points analyzed (X and Y values) is also provided. These data highlight variations in dabrafenib sensitivity across different deletion lines, supporting insights into how SOX10 enhancer loss influences drug response.

**Supplementary Table 5. IC50 and dose-response data for WM115 and A375 cell lines treated with trametinib.**

| **WM115 Trametinib** | | | | | | |
|---|---|---|---|---|---|---|
| | W_WT | W_1.1 | W_1.2 | W_1.3 | W_4.1 | W_4.2 |
| **log(inhibitor) vs. normalized r esponse** | | | | | | |
| **Best-fit values** | | | | | | |
| **LogIC50** | -0.8379 | -1.019 | 0.8272 | -0.401 | 0.5894 | 0.07334 |
| **IC50** | 0.1453 | 0.09582 | 6.718 | 0.3972 | 3.885 | 1.184 |
| **95% CI (profile likelihood)** | | | | | | |
| **LogIC50** | -1.138 to -0.5265 | -1.396 to -0.6576 | 0.2689 to 1.341 | -0.8014 to -0.01499 | -0.0007598 to 1.214 | -0.4193 to 0.7153 |
| **IC50** | 0.07277 to 0.2975 | 0.04019 to 0.2200 | 1.857 to 21.92 | 0.1580 to 0.9661 | 0.9983 to 16.38 | 0.3808 to 5.191 |
| **Goodness of Fit** | | | | | | |
| **Degrees of Freedom** | 17 | 17 | 17 | 17 | 17 | 17 |
| **R squared** | 0.832 | 0.7504 | 0.6759 | 0.7768 | 0.5407 | 0.4484 |
| **Sum of Squares** | 3353 | 4404 | 5829 | 3896 | 7216 | 8565 |
| **Sy.x** | 14.04 | 16.1 | 18.52 | 15.14 | 20.6 | 22.45 |
| **Number of points** | | | | | | |
| **# of X values** | 18 | 18 | 18 | 18 | 18 | 18 |
| **# Y values analyzed** | 18 | 18 | 18 | 18 | 18 | 18 |

| **A375 Trametinib** | | | | | |
|---|---|---|---|---|---|
| | A_WT | A_1.1 | A_1.2 | A_4.1 | A_4.2 |
| **log(inhibitor) vs. normalized response** | | | | | |
| **Best-fit values** | | | | | |
| **LogIC50** | -0.08063 | -0.00558 | 0.6316 | 0.3189 | 0.7064 |

| | | | | | |
|---|---|---|---|---|---|
| **IC50** | 0.8306 | 0.9872 | 4.281 | 2.084 | 5.086 |
| **95% CI (profile likelihood)** | | | | | |
| **LogIC50** | -0.3811 to 0.2301 | -0.2804 to 0.2816 | 0.02353 to 1.232 | -0.1557 to 0.9313 | 0.09666 to 1.390 |
| **IC50** | 0.4158 to 1.699 | 0.5243 to 1.913 | 1.056 to 17.05 | 0.6988 to 8.537 | 1.249 to 24.54 |
| **Goodness of Fit** | | | | | |
| **Degrees of Freedom** | 17 | 17 | 17 | 17 | 17 |
| **R squared** | 0.8145 | 0.8507 | 0.4939 | 0.6159 | 0.5949 |
| **Sum of Squares** | 3696 | 3023 | 7241 | 8273 | 12940 |
| **Sy.x** | 14.75 | 13.34 | 20.64 | 22.06 | 27.59 |
| **Number of points** | | | | | |
| **# of X values** | 18 | 18 | 18 | 18 | 18 |
| **# Y values analyzed** | 18 | 18 | 18 | 18 | 18 |

**Supplementary Table 5. IC50 and dose-response data for WM115 and A375 cell lines treated with trametinib.**
This table provides the results of dose-response experiments for WM115 and A375 cell lines, including wild-type (WT) and SOX10 enhancer deletion lines, treated with the MEK inhibitor trametinib. Best-fit values for log(IC50) and IC50 are reported alongside 95% confidence intervals (CI). Goodness-of-fit metrics include R-squared values, degrees of freedom, and sums of squares. The total number of data points analyzed (X and Y values) is also provided. These data highlight variations in trametinib sensitivity across different deletion lines, supporting insights into how SOX10 enhancer loss influences drug response.

**Supplementary Table 6. Correlation matrix for phenotype scores, IC50 values, and gene expression levels.**

| | Score | Score Shift | Dab IC50 | Tram IC50 | NTRK1 expression | NTRK2 expression | NTRK3 expression | SOX10 expression |
|---|---|---|---|---|---|---|---|---|
| **Score** | 1 | 0.405419 | 0.292734 | 0.603918 | 0.714177 | -0.457515 | 0.104775 | -0.085682 |
| **Score Shift** | 0.405419 | 1 | 0.588625 | 0.614266 | -0.158844 | 0.442559 | -0.489333 | -0.662599 |
| **Dab IC50** | 0.292734 | 0.588625 | 1 | 0.787869 | -0.059974 | 0.140497 | 0.012772 | -0.871855 |
| **Tram IC50** | 0.603918 | 0.614266 | 0.787869 | 1 | 0.29034 | 0.091727 | -0.098578 | -0.633561 |
| **NTRK1 expression** | 0.714177 | -0.158844 | -0.059974 | 0.29034 | 1 | -0.743404 | 0.586583 | 0.439693 |
| **NTRK2 expression** | -0.457515 | 0.442559 | 0.140497 | 0.091727 | -0.743404 | 1 | -0.655727 | -0.501215 |
| **NTRK3 expression** | 0.104775 | -0.489333 | 0.012772 | -0.098578 | 0.586583 | -0.655727 | 1 | 0.399253 |
| **SOX10 expression** | -0.085682 | -0.662599 | -0.871855 | -0.633561 | 0.439693 | -0.501215 | 0.399253 | 1 |

**Supplementary Table 6. Correlation matrix for phenotype scores, IC50 values, and gene expression levels:** ››Pearson correlation coefficients between phenotype scores (Score), score shifts, IC50 values for dabrafenib (Dab IC50) and trametinib (Tram IC50), and expression levels of key genes (*NTRK1*, *NTRK2*, *NTRK3*, and *SOX10*). Positive and negative correlations indicate relationships between these variables across the cell lines studied. Strong correlations are highlighted, providing insights into the interplay between phenotype shifts, drug sensitivity, and gene expression in SOX10 enhancer-deleted lines.

**Supplementary Table 7. HiFi primers for creating reporter construct vectors targeting Regions 1 and 4.**

| *Region* | *Direction* | *Sequence* |
|---|---|---|
| *Region 1* | Forward | atggtccagcctgcttttttgtacaaacttgggCTTTAGCCATGATGCCCTCTGACCTTT |
| | Reverse | cagagactccctggtgtctgaaacacaggccagatgggccCGGCACCCCGGC |
| *Region 4* | Forward | attggccatggtccagcctgcttttttgtacaaacttgggGAGCTTGGAGCAAGCAGCT |
| | Reverse | cagagactccctggtgtctgaaacacaggccagatgggccCGCCGCGGCCGC |

**Supplementary Table 7. HiFi primers for creating reporter construct vectors driven by Regions 1 and 4.**
Forward and reverse HiFi primers used for cloning Regions 1 and 4 into reporter construct vectors. Primers were designed to amplify the regions of interest with blunt ends for NEBuilder HiFi DNA Assembly. The sequences include overlaps tailored for vector integration, enabling the generation of constructs for functional analysis of enhancer activity. The direction (forward or reverse) and specific sequences for each region are indicated. Adjustments to primer length were made to meet synthesis constraints.

**Supplementary Table 8. gRNA sequences for CRISPR/Cas9-mediated deletion of Regions 1 and 4.**

| | **Genomic Coordinates** | **Sequence** |
|---|---|---|
| **Region 1:** | chr22:38412487-38412987 (hg19) | CTTTAGCCATGATGCCCTCTGACCTTTCATAAATCAGAGGGGCTTAGGGGCGAGGGGG<br>CTGCTTGGCAGGACTTGGTGGGGTGGGGGCTTAGAAGCAGCTGCGCGCGACGTTGAC<br>ATTGTTCCCACCATTCTAAGTGCAACAAATCCCTCTATTGTGTTCCTGGTTTATCTGGTT<br>CCTCTTGTTTATGAGCAGGGCTCCTTTGGCAGCGGTTCCAGCCCTGGGCGGGTGGAA<br>AGAGTGCTGGCACGCACCGCGGGGGGAGGGGGCGGGAGGGGGCCGCCGTCAATGCC<br>CGcattgtccccgcgctttttgtTTCTACTGTAATGACATGTTGGAATAGAGAGAGAGAGAGAAGG<br>AAAAAAAAAAAGATACAAGCTGGGGAGGGAAGAGGAGGGCAGAAAAGAAAGAGCA<br>GAGCAAGGGCCTGGTGTGGATTGATGTCTGGGCCTGGGCGGATGTGCCGGCAGCTGG<br>GCGGCCGGGAGAGATGGAGCCAGGCCGGGGTGCCG |
| | gRNA1 | GTGCCGGGGGGCAGCGCAAGNGG |
| | gRNA2 | AGCTGCCGGCACATCCGCCCNGG |
| | gRNA3 | AAGGGACCATTGTGTCCGAANGG |
| | gRNA4 | TCTGACCTTTCATAAATCAGNGG |
| **Region 4:** | chr22:38434349-38434849 (hg19) | GAGCTTGGAGCAAGCAGCTTGGGGGAAAGGACCAGGCCGGGCCCAGGCAGGTGGA<br>AGCCATGGCTCTGGCCGGGTCACAGGGCGGCTGGGGTTTGACCTGGCGGCCATGATG<br>GGCTCTGGACCCAGGAGCCACCCGGGCACAGGATGGGACGGGTTGAGAGATTGGTC<br>GTCCAGGCGTTGAGTGTGCCCTCGCTCCTCCCCACCTTCCGCCGTGGCCTTGAGCCCC<br>TTGCGCCCGCGCCCATGGGGTTAAATCTCTCCCTGTCTCTCTCTGCTCCAAGTTGTTTT<br>CCAAGTGAGGCAGAGAATGGTTCTCTTGTTACCAACATGGCTTCTGGGCATTGGGTA<br>ATGCGCTCCCTCTTCTCCCGCAGCGCCacaaagggaccttttgtTCCCCTCCCTGCCCCTCTCGC<br>TGGCTCTTCCCGGCCCCCCACCCACCCAAACTCCCTCCTCCCCCCGCCGGCGCCCGCA<br>CCCCGGGGCTGCGCGCTGACCGTGGCGGCCGCGGCG |
| | gRNA1 | ACGGTCAGCGCGCAGCCCCGNGG |
| | gRNA2 | TCGGGGCCGGGGCAGTCGTANGG |
| | gRNA3 | GAGGGCGTCCTCTTGAGGTTNGG |
| | gRNA4 | ATGGCTCTGGCCGGGTCACANGG |

**Supplementary Table 8. gRNA sequences for CRISPR/Cas9-mediated deletion of Regions 1 and 4.**
Guide RNA (gRNA) sequences designed for CRISPR/Cas9-mediated deletion of Regions 1 and 4 upstream of *SOX10*. Each region

includes the genomic coordinates (hg19), target sequences, and corresponding gRNAs. The gRNA sequences were optimized to target specific sites within the regions of interest while minimizing off-target effects.
